# Real-time tracking of drug binding to Influenza A M2 reveals a high energy barrier

**DOI:** 10.1101/2023.04.07.536045

**Authors:** Kumar Tekwani Movellan, Melanie Wegstroth, Kerstin Overkamp, Andrei Leonov, Stefan Becker, Loren B. Andreas

**Author notes:** **Corresponding Author:** Loren B. Andreas, **Email:**. **Author Contributions:** Kumar Tekwani Movellan performed the experiments. Stefan Becker, Melanie Wegstroth, Kerstin Overkamp and Andrei Leonov expressed, purified and reconstituted the protein samples. Kumar Tekwani Movellan and Loren B. Andreas designed the experiments and wrote the manuscript. All authors have given approval of the manuscript. **Competing Interest Statement:** No competing interests.

## Abstract

The drug Rimantadine binds to two different sites in the M2 protein from influenza A, a peripheral site and a pore site that is the primary site of efficacy. It remained enigmatic that pore binding did not occur in certain detergent micelles, and in particular incomplete binding was observed in a mixture of lipids selected to match the viral membrane. Here we show that two effects are responsible, namely changes in the protein upon pore binding that prevented detergent solubilization, and slow binding kinetics in the lipid samples. Using 55-100 kHz magic-angle spinning NMR, we characterize kinetics of drug binding in three different lipid environments: DPhPC, DPhPC with cholesterol and viral mimetic membrane lipid bilayers. Slow pharmacological binding kinetics allowed the characterization of spectral changes associated with non-specific binding to the protein periphery in the kinetically trapped pore-apo state. Resonance assignments were determined from a set of proton-detected 3D spectra. Chemical shift changes associated with functional binding in the pore of M2 were tracked in real time in order to estimate the activation energy. The binding kinetics are affected by pH and the lipid environment and in particular cholesterol. We found that the imidazole-imidazole hydrogen bond at residue histidine 37 is a stable feature of the protein across several lipid compositions. Pore binding breaks the imidazole-imidazole hydrogen bond and limits solubilization in DHPC detergent.

**Highlights:** 5 °C kinetically traps Influenza A M2 (as dimer of dimers) in the apo form *in vitro*.

Kinetic control allows characterization of non-specific chemical shift perturbation.

Pore-bound M2 loses dissolvability in DHPC micelles, suggesting structural change.

M2, residues 18-60, forms a dimer-of-dimers structure in several bilayer compositions.

## Introduction

The tetrameric matrix protein 2 (M2) from influenza A(1-3) forms a central pore through which protons can permeate at low pH. The protein is the target of two aminoadamantyl drugs amantadine and rimantadine that bind to the pore.(4-6) These drugs alter the pH sensitive proton conduction of the protein, and thereby interfere with the endosomal infection pathway in which low pH triggers release of viral RNA.(3, 7) Widespread resistance to these inhibitors in recent flu seasons has led to removal of both inhibitors from the market, and replacements are sought for the common rimantadine resistant variants. These efforts have led to the identification of several compounds that block resistant strains.(8-12)

The full length protein is 97 residues, and each monomer contains a largely unstructured N-terminus, which points towards the outside of the virus particle, a single pass transmembrane (TM) alpha helix followed by an amphipathic helix, and finally a C-terminal domain proposed to interact with matrix protein 1.(13) The minimum sequence needed to recapitulate the conduction and drug inhibition properties of the full length protein is comprised by the TM and amphipathic helices, roughly residues 18-60 or 22-62.(14-16) Such protein constructs are referred to as the conductance domain (CD). The TM domain alone is still drug sensitive, and its structure has been captured in a variety of conformations, both with(4, 6, 17) and without drug,(18) and at a resolution allowing the measurement of precise water positions.(10, 19-21)

Structures have been reported for CD constructs in both detergent and lipid environments. (16, 17, 22-25) Of these, fourfold symmetric structures were determined from a detergent environment by solution NMR, as well as a crystal structure of V27A M2 in lipidic cubic phase in complex with a spiro-adamantane molecule.(17) While the first crystal structure of M2 using the TM M2 peptide, residues 22 to 46 was asymmetric,(6) later structures of TM M2, including structures solved in lipidic cubic phase exhibited 4-fould symmetry.(18, 19, 21) In contrast, two-fold symmetry has been observed in lipid bilayer conditions using CD M2, for both 1,2-diphytanoyl-sn-*glycero*-3-hosphatidylcholine (DPhPC) and 1-palmitoyl-2-oleoyl-*sn*-glycero-3-phosphocholine (POPC).(26) In this case, two distinct sets of peaks are observed by MAS NMR, which when assigned result in two distinct sets of resonances. We refer to these as chain A and chain B. The twofold dimer-of-dimers symmetry is observed for many residues of M2_18-60_, and is particularly evident at H37, which forms intermolecular imidazole-imidazole hydrogen bonds.(27) Spectra of CD M2 in more complex lipids or in 1,2-dimyristoyl-sn-glycero-3-phosphocholine(DMPC) demonstrate broader lines, and a dimer-of-dimers structure was not detected under these conditions for M2_21-61_. (28)

In the earliest work on oriented TM domain samples, conformational changes were detected upon addition of amantadine that allowed a detailed determination of the monomer structure of the bound form.(29) Later measurements on the CD domain succeeded in determination of the apo structure of both the transmembrane and amphipathic helices, from which a tetrameric structure was assembled based on imidazole-imidazolium dimerization.(25, 30) Intriguingly, a chimeric protein containing the drug binding site from influenza A M2, but the C-terminal sequence of influenza B M2, showed changes in helix packing upon rimantadine binding,(5) yet the influenza A tetramer in DHPC micelles was insensitive to the drug.(24) In all such cases where the drug binds to the pore, large chemical shift changes, exceeding 1 ppm in both ^15^N and ^13^C dimensions, are also observed throughout the protein, suggesting again that the drug induces a repacking of the transmembrane helices. The large shift changes associated with pore binding are shown in Figure 1 for rimantadine binding to CD M2 (residues 18-60) reconstituted in a DPhPC lipid bilayer.

**Figure 1.**
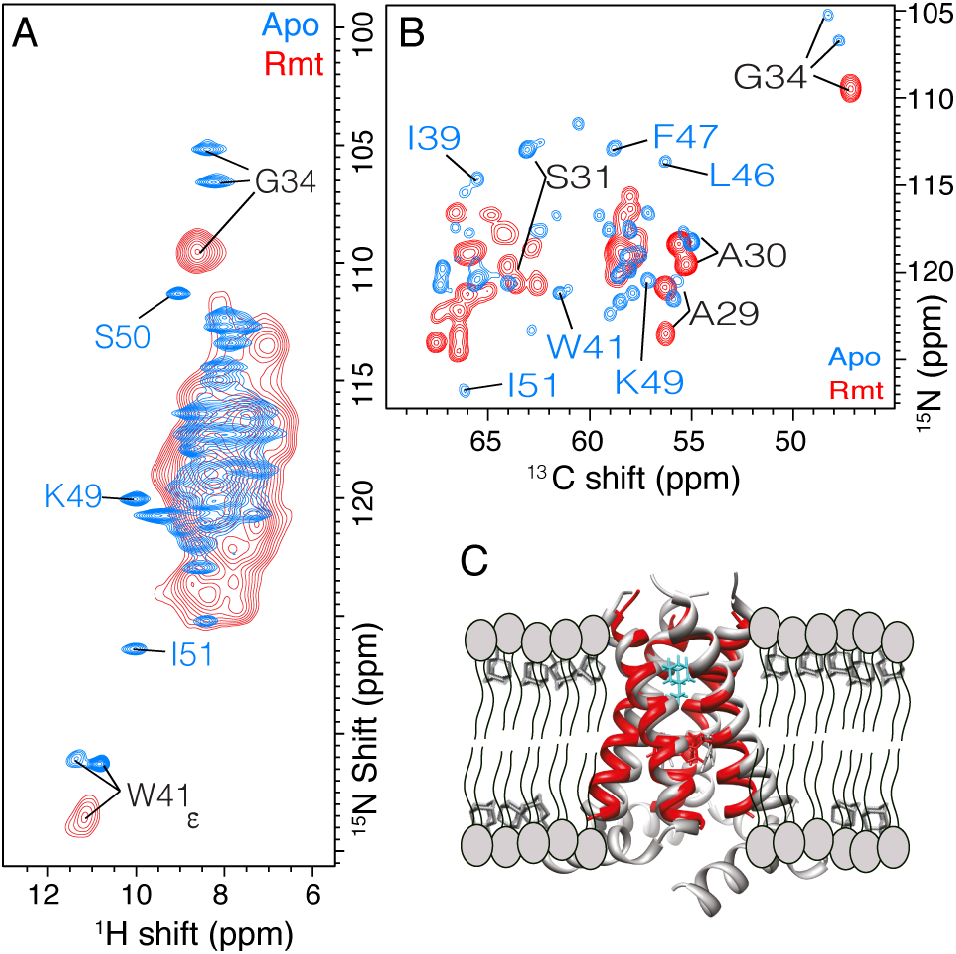
Chemical shift changes induced by pore binding with 40 mM Rmt. In A), 2D (H)NH spectra of M2 show the large changes between apo (blue), and specific binding (red) of the inhibitor rimantadine (Rmt). In B), the ^13^C^15^N projections of (H)CANH spectra show chemical shift perturbation of the Cα and N^H^ backbone resonances. A large excess of 16 molecules of Rmt drug was used per tetramer. In C) the pore binding location (cyan) is shown on the CD model, pdb 2L0J in grey and on the bound TM structure 6BKL in red. The spectra were recorded in a 950 MHz Bruker spectrometer equipped with a 0.7 mm 4 channel probe at 100 kHz MAS with cooling gas set to 260 K (sample temperature of approximately 10 to 15 °C).

Despite this wealth of structural data in the literature, it remains unexplained how the functionally relevant CD did not bind drug in the pore when reconstituted in DHPC micelles.(24) Similarly, the CD construct only partly bound drug to the pore when reconstituted in lipid bilayers with concentrations of cholesterol and sphingomyelin selected to match the composition in viral membranes, yet complete binding was observed for the TM domain in these same lipids,(28) and also for the CD in 1,2-diphytanoyl-sn-*glycero*-3-hosphatidylcholine (DPhPC) lipids.(26)

Here we investigate the potential reasons behind these observations via NMR. We used M2_18-60_ reconstituted in lipids as a dimer-of-dimers structure and characterized two effects. Firstly, we find that slow kinetics of drug binding, which are modulated by the lipid composition and pH, can kinetically trap the sample in the pore-unbound state. Secondly, the protein solubility in detergent is altered when drug is bound to the pore. Control of the kinetics via low temperature additionally allows separation of specific and non-specific chemical shift perturbation (CSP), and to associate the large changes in chemical shift at functionally important residues to the specific effects of the drug.

## Materials and Methods

Expression and lipid reconstitution was carried out as described previously.(22, 24, 26, 31) The conductance domain (CD), which comprises residue 18 to 60, of Influenza A was used, taking the sequence of the M2 Udorn/72 H3N2 strain. The Udorn sequence has often been referred to as wild type (WT) and has serine at position 31. Cysteines 19 and 50 were replaced by serine, as in previous studies. The resulting amino acid sequence is RSNDSSDPLVVAASIIGILHLILWILDRLFFKSIYRFFEHGLK. Briefly, the expression was performed in *E. coli* (BL21DE3) using an N-terminal TrpLE fusion to the above Udorn sequence. The protein was purified using standardized purification used for inclusion bodies. The inclusion body pellet was resuspended in a solution of 50 mM sodium phosphate, 6 M guanidine at pH 6.8. After centrifugation the supernatant was passed through a nickel affinity column (Ni-NTA), and eluted in 50 mM sodium phosphate buffer containing 400 mM imidazole at pH 6.8. Elution fractions were collected and dialyzed against water. The white precipitated was dissolved in 70 percent formic acid and cleaved with an excess of cyanide bromide. After cleavage, the peptide was lyophilized, dissolved in 2:1:1 hexafluoroisopropanol: formic acid: water and finally passed through a C4 reverse phase HPLC column using a gradient with elution in isopropanol/acetonitrile/water. The pure protein was refolded in a 2% solution of Octyl-Beta-glucoside (in 40 mM sodium phosphate, 30 mM glutamate, 3 mM sodium azide) and then reconstituted in perdeuterated lipids (95% d78-phytanoyl 50% d at the alpha position, methyl-d9-choline) 1,2-diphytanoyl-sn-glycero-3-phosphatidylcholine lipids (DPhPC, FBreagents) using a lipid to protein ratio (LPR) of 1 to 1 by mass (25 to 1 by mole of tetramer) or in DPhPC with 30% cholesterol using an LPR of 1.18 to 1 by mass or a mix of 1,2-dipalmitoyl-sn-glycero-3-phosphocholine (DPPC), 1,2-dipalmitoyl-sn-glycero-3-phosphoethanolamine (DPPE), egg SM Sphingomyelin (from Avanti) and cholesterol referred to as viral membranes (VM) with a molar ratio of 21:21:28:30 (%) using an LPR of 1.24 to 1 by mass. In each case, detergent was slowly removed via dialysis against NMR buffer (30 mM glutamate, 40 mM phosphate, 0.02% sodium azide, pH 7.8) for reconstitution and resulted in the appearance of a white precipitate containing protein and lipids. To ensure similar protein amounts among the different samples, the precipitate was concentrated into a membrane pellet via centrifugation, and prepacked into a 1.3 mm Bruker rotor. Either the pellet from a single rotor, or freshly reconstituted protein, was then pushed into a 200 μL solution of 40 mM Rmt at 4 °C and left overnight before the sample was packed back into the rotor. We estimated the protein concentration based on the mass and the volume inside of a 1.3 mm rotor, volume 3-4 μL and a protein mass of 1-1.5 mg. Thus, we estimate a protein tetramer concentration of around 4 to 6.5 mM. An additional sample for each of the three lipid compositions was prepared using a 20 to 1 lipid to protein mass ratio (about 450 lipids per tetramer). For a single sample prepared at higher drug concentration, 0.2 mg Rmt was added directly to the membrane pellet.

To track the kinetics of drug binding, the packed rotor was incubated at different controlled temperatures. An Eppendorf with 500 μL of water was pre-warming to the set temperature in a water bath. The rotor was then transferred from ∼4 °C to the prewarmed water and incubated for the required time. After incubation, the rotor was transferred to an Eppendorf tube containing 500 μL of water at ∼4 °C. The rotor was then transferred to an 800 MHz Bruker spectrometer for (H)NH acquisition. In order to avoid conversion during acquisition of the data, we kept the temperature as low as possible by increasing the gas flow to 1200 l/h with a set temperature of 240 K and increasing the spinning rate slowly up to 40 kHz. This ensures a sample temperature of about 5 °C. Following acquisition of the (H)NH spectrum, which took only several hours, the sample spinning was stopped, and the rotor was returned to ∼4 °C in preparation for the next incubation step, repeating the process. The binding of Rmt was monitored by the disappearance of characteristic apo state peaks, G34 NH and H37 side chain. The binding can also be tracked via the appearance of characteristic pore-bound peaks, which are still resolved in the 40 kHz MAS (H)NH spectrum for G34.

For tests of detergent solubilization, the unbound and pore bound protein from the 1.3 mm rotors were resuspended in a 300 mM DHPC solution, which is identical to the detergent conditions previously reported for the solution NMR structure determination.(24) These samples were centrifuged at low speed (16,000 g with a benchtop centrifuge) and the supernatant was transferred to a 3 mm NMR solution tube and measured in a 600 MHz Bruker NMR spectrometer equipped with a cryoprobe. The pellet from the resolubilized pore-bound sample was packed in a 1.3 mm rotor and measured at in 800 MHz Bruker NMR spectrometer equipped with a three-channel narrow bore 1.3 mm probe at 55 kHz MAS.

NMR data were processed using Bruker Topspin 3.6 and analyzed using CcpNmr.(32)

In order to estimate the activation energy, we assumed Arrhenius behavior and estimated the initial rate as the slope between about 95% and 50 % of the initial intensity (see table S6 for exact information about the values used for the plots). The error estimation includes only the spectrum noise, and is therefore a minimum error estimate. It does not include other potential sources of error, for example in the timing (or temperature) of the incubation step, which may arise due to accidental warming of the sample during handling.

Assignment data was acquired at a 950 MHz Bruker NMR spectrometer with a 0.7 mm narrow bore MAS NMR probe using amide proton detected assignment spectra.(33) The measurement was acquired using VT gas set to a thermocouple temperature of 260 K (sample temperature of approximately 10 to 15 °C) and with 100 kHz MAS for apo M2. For non-specific binding, the MAS was reduced to 80 kHz to reach a sample temperature of about 5 degrees, which prevents drug binding during the measurement.

## Results and Discussion

### Specific rimantadine binding in the pore

Recombinant M2_18-60_ was reconstituted in perdeuterated DPhPC lipids. As previously described, the M2 tetramer assembles as a dimer-of-dimers structure under these conditions.(22, 26, 27, 34) The sample was incubated with 40 mM rimantadine (Rmt) and then packed into a 1.3 mm MAS NMR rotor. After rotor packing, the sample was placed in a water bath to control incubation time and temperature during binding. Incubation was paused by transferring the rotor to a 4 °C water bath and the measurement at the spectrometer was carried out at low enough temperature that minimal binding occurs during measurement. Figure 1 shows proton detected magic-angle spinning (MAS) nuclear magnetic resonance (NMR) spectra of an apo sample (Figure 1A, blue), as well as a Rmt-containing sample, which has fully converted to the bound state (Figure 1A, red). Large chemical shift perturbations are evident, and they occur due to binding of Rmt to the pore as previously observed.(26, 28) Note in particular the ∼7 ppm change of the S31 amide nitrogen (shown in the ^13^C^15^N projection of the 3D (H)CANH spectrum, Figure 1B). Consistent with previous studies in phosphocholine lipid bilayers, large chemical shift perturbation (CSP) of > 2 ppm for ^15^N and > 1 ppm for ^13^C, occur throughout the TM region residues 27 to 42 upon pore binding.(22, 28, 35) We refer to pore binding as specific since it is the primary site of inhibition. An increase in linewidths is observed in the (H)NH spectra of the pore-bound sample (Figure 1A, red). This likely results from changes in structural dynamics, but might also reflect changes in helical packing within the tetramer structure of M2 upon pore binding. In the (H)CANH spectrum (Figure 1B, red) most of the peak doubling is no longer resolved, due in part to a smaller difference between the chemical shifts of A and B peaks, which is suggestive of a more symmetric structure upon pore binding. These interpretations are also consistent with the loss of the interhelical hydrogen bond at residue 37 in the rmt-bound state, which was previously reported.(27) As in previous studies, when the sample was handled at room temperature, rather than 4 °C, complete binding was observed.

To further characterize the bound state, which also exhibits peak doubling, a full set of proton-detected 3D ultra-fast MAS NMR measurements were acquired for assignment proposes.(33) We were able to identify two distinct set of peaks for residues V26 to G35 in the bound state. We name these as chain A and B. Inter-residue linking spectra connecting residues A30 to I35 of both A and B are shown in Figure S12 and S13. Note that for this pore bound case, it is not known if the two chains assemble as one tetramer or come from distinct tetramer conformations. Unambiguous assignments span from residue L26 to I35 for Chain A and from L26 to I42 for Chain B (Table S3). Weak or missing cross-peaks for residues L36 and H37 of Chain A prevented unambiguous assignment of these residues. Based on the assignments, we determined chemical shift perturbations of the pore bound state compared to the apo for individual residues including H^N^, N and Cα shifts (Figure 2A, S14). Note that there are two possible comparisons due to two sets of peaks for both apo and bound states. Both comparisons yield qualitatively similar results with large CSP for pore binding.

**Figure 2.**
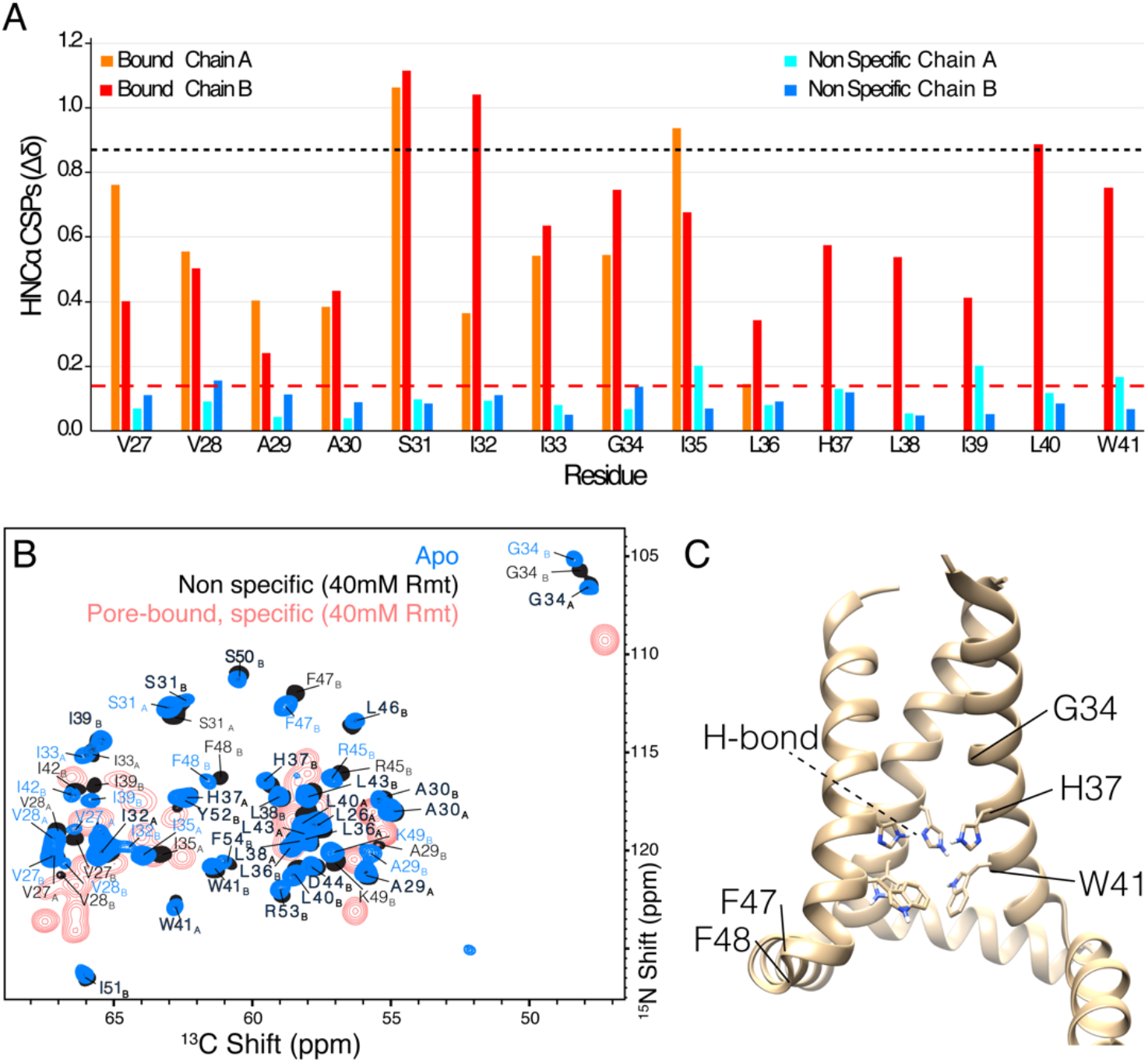
Non-specific CSP (blue shades) compared with specific CSP (red and orange) for the chains A and B as indicated in the legend. In A) the CSP is shown per residue by combining H^N^, Cα, and N chemical shifts as described in the text. The dashed red line shows the CSP value that is 3 times the root mean square of non-specific CSP (∼0.14 ppm). The CN projections of the 3D (H)CANH spectra are shown in B) with apo in blue and non-specific in black. Subscripts indicate the chain, either A, or B. The spectrum of pore-bound M2 from Figure 1 is reproduced in red for comparison. In C) the tetrameric arrangement of M2 is shown, based on the structure of pdb 2L0J. Note that A and B chains in 2L0J have nearly the same backbone structure and the symmetry is broken only in the hydrogen bonding arrangement of residue H37 (indicated by the dashed black line). Selected residue positions are indicated: the pH sensor H37, the gating residue W41, G34 in the pore, and F47-48 at the junction between transmembrane and amphipathic helices. Spectra were acquired at a 950 MHz Bruker NMR spectrometer with a 0.7 mm narrow bore MAS NMR probe. The VT gas was set to a thermocouple temperature of 260 K (sample temperature of approximately 10 to 15 °C) and with 100 kHz MAS for apo M2. For non-specific, the MAS was reduced to 80 kHz to reach a sample temperature of about 5 degrees, which prevents drug binding during the measurement.

### Characterization of non-specific rimantadine binding

In contrast, when the sample is kept at 4 °C, only small chemical shift changes, below 0.5 ppm, were observed in the presence of drug (Figure 2). We refer to these small changes as non-specific effects of the drug. These non-specific shift changes can be compared with the pore-bound shift changes in Figure S1A-D. Even after several weeks of incubation at 4 °C, peaks indicative of pore binding were absent, while both pore-bound and non-specific (i.e. pore-unbound) populations were present after a week of incubation at 20 °C (Figure S1D-E). The two populations, non-specific and specific resonance positions, can be seen clearly for G34 (Figure S1E), with distinct ^15^N chemical shifts at about 106.5 ppm and at 109.5 ppm, respectively. The chain B resonances in the apo state could be assigned continuously up to residue 54, while the chain A resonances were only assigned to residue 43. It is unclear whether this difference is due to structural dynamics in Chain A or rather that the Chain A and Chain B chemical shifts could not be distinguished for C-terminal residues. Note that two additional residues (53-54) could be assigned as compared with our previous ^13^C-detected assignments from S31N M2.(22) The slow kinetics of pore binding at 4 °C is qualitatively consistent with the fact that glycerol was observed to kinetically prevent pore binding.(31) The glycerol was used in these previous studies in order to form a glassy sample for low temperature dynamic nuclear polarization (DNP) enhanced measurements.

Kinetic control allowed characterization of non-specific CSP. Apo assignments were carried out at 100 kHz (Figure S2-S3). For assignment of the non-specifically drug-bound sample, we reduced the spinning to 80 kHz MAS with a 0.7 mm rotor and decreased the set temperature to 250 K to reach a sample temperature of ∼5 °C to minimize pore binding by Rmt during acquisition. Figure 2B shows the excellent resolution obtained under these conditions for a non-deuterated sample, which allowed us to transfer apo assignments to the non-specific bound state using 3D triple resonance spectra and obtain residue specific CSP information. Consistent with previous reports,(22, 26, 36) the existence of two sets of peaks in the TM region indicates a dimer of dimers structure. The two sets of peaks are indexed here as chain A (for which assignments extend from L26 to W41), and chain B (assignments from residue V27 to F54). P25 was not assigned here, although it is part of the rigid helix,(26) since it is not detected in amide proton based spectra. The CSP was calculated for each residue as 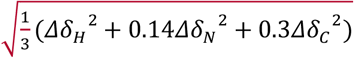,(37) and the individual non-specific CSP for each of H^N^, N, and Cα, is shown in Figure S4. The chemical shifts are tabulated in Tables S1-S3.

All non-specific CSP values are small, similar to CSP observed previously between two lipid environments.(26) In the TM region, chain A shows elevated non-specific CSP, above 3 times the standard deviation (> ∼ 0.14 ppm), toward the C-terminus, from G34 to W41 with a cluster of larger values of ∼0.2, ∼0.2 ppm and ∼0.17 for residues I35, I39 and W41 respectively. In contrast, chain B shows elevated non-specific CSP for residues V28, ∼0.16 ppm, and G34, 0.15 ppm, located at the N-terminus of the TM. The ∼0.1 to 0.2 ppm non-specific CSPs are much smaller than the averaged CSPs observed for specific binding to the pore, which are nearly 1 ppm for several residues spanning the TM helix, and that have previously been attributed to conformational changes.(26, 35, 38) The small non-specific CSPs are likely due to a combination of partitioning of drug to the membrane(28, 39) as well as to association at the peripheral binding site identified previously,(24) and consistent with DNP data.(31) Elevated Cα non-specific CSP of above 0.35 ppm was observed for residues F47, F48, and Y52 near this peripheral site (Figure S4). The non-specific CSP of ^15^N, ^13^C and ^1^H are plotted individually in Figure S4. While the CSPs for non-specific binding are small in comparison to specific CSPs linked to changes in the tetramer quaternary structure,(27) they are distributed across the helices and include the pore facing residue G34, which is sensitive to membrane conditions and can form a kink.(38) This emphasizes a small but detectable degree of sensitivity of the channel to the change in environment, a topic explored in more generally for the TM construct.(40)

Residues H37 and W41 are key functional residues controlling conduction in response to pH. It has recently been shown that there is an imidazole - imidazole hydrogen bond in the Apo state and that the hydrogen bond is broken upon Rmt pore binding.(27) H37 N- -H–N hydrogen bonding persists at a lower pH of 6.2 in a full length construct.(41) Non-specific binding causes small CSP at the H37Nε and the imidazole - imidazole hydrogen bond was still present (Figure 3). This indicates that the non-specific effects and peripheral binding likely have at most a minor influence on the pore structure. However, since the pore-bound preparations are also likely bound peripherally, we cannot exclude that the two binding locations act cooperatively.

**Figure 3.**
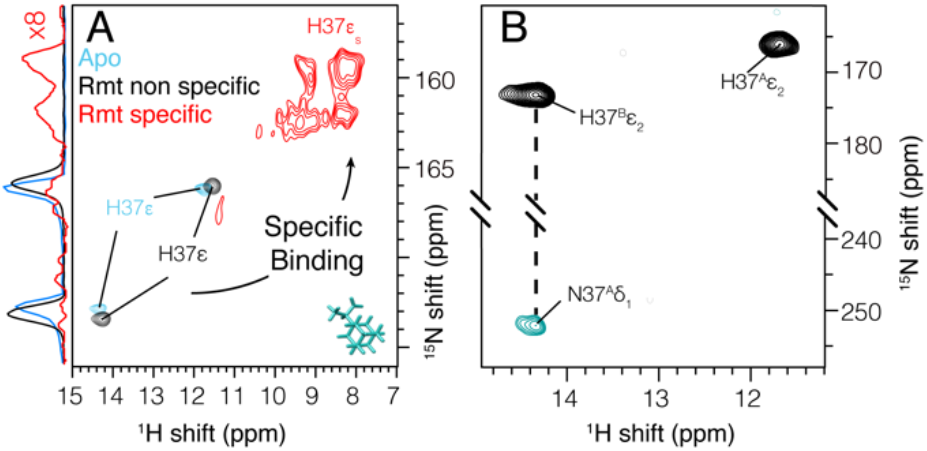
Histidine side-chain CSP and hydrogen bonding. In A) (H)NH spectra show non-specific changes (black) that are small compared with specific pore binding (red). In B), the homonuclear nitrogen J-coupling was evolved for 36 ms, and the negative peak (blue-green) is indicative of the imidazole-imidazole hydrogen bond. Data were recorded at a 950 MHz spectrometer. Spectra were recorded at 950 MHz using 80 kHz MAS and a 0.7 mm rotor to maintain a low sample temperature.

### Measurements of Rimantadine pore-binding kinetics in different environments

To kinetically characterize the binding process, samples were spun at reduced rates in order to keep the temperature of the sample at ∼ 5 °C during NMR measurement. For 1.3 mm rotors, this was only 40 kHz MAS with cooling gas at 240 K. We primarily used low LPR samples of 1 to 1 by mass, which correspond to about 25 lipids per tetramer. This lipid composition was optimized previously from an initial condition with about 50 lipids per tetramer, which resulted in an identical spectrum.(26) These conditions result in an acquisition time of only a few hours for the (H)NH spectrum, which reduces the potential risk of pore binding during acquisition. To control the drug concentration, about 1 mg of M2 protein (∼3-4 μL) was pre-packed in a 1.3 mm rotor and incubated in 200 μL of 40 mM Rmt at 4 °C overnight. The samples, then in the non-specific bound state, were re-packed in either 0.7 mm or 1.3 mm rotors. After recording an initial (H)NH spectrum, the entire packed rotor was then incubated at 25, 40, or 55 °C in a water bath. The rotors were kept closed during the incubation process, which ensures consistent sample conditions at each time point. Binding was tracked by periodically removing the rotor from the water bath, recording an (H)NH spectrum, and replacing the sample in the bath. For measurement conditions of 40 kHz MAS with a non-deuterated membrane protein sample, the resolution is not sufficient to characterize binding kinetics using signals from all amino acids. Residues that report on Rmt pore binding and are particularly well resolved in the 2D spectrum are the side chain of H37 and the backbone of G34. We therefore tracked the intensities of H37Nε2 and G34N of both chains over time and at different temperatures (Figure 4 and Figure S5), and used these data to estimate the energy barrier of Rmt binding (Figure S6 and Table S7). At a 55 °C incubation temperature, full binding is observed in a few hours, while at 40 °C, binding occurred within 3 days. Figure 4 shows the combined intensities of H37Nε2 from chains A and B. Surprisingly, at temperatures of 25 °C (Figure 4, blue) and 40 °C (Figure 4, green) the curves do not follow an exponential decay as would be expected for pseudo first order kinetics but rather proceed more slowly at first, indicating kinetic cooperativity. Expectation of pseudo first order kinetics justified by drug concentration in high excess. Note also that Rmt partitions to the membrane(28, 39) A possible explanation for non-exponentiality is that neighboring tetramers may interact in the dense and highly concentrated NMR sample, such that binding is accelerated when a neighboring tetramer is already bound. This is consistent with the recent observation by MAS NMR and coarse-grained molecular dynamics simulations showing that M2 has a tendency to cluster in lipid bilayers.(42) The slow kinetics is in contrast to fast blocking of proton currents observed in functional assays as discussed further below.(16)

**Figure 4.**
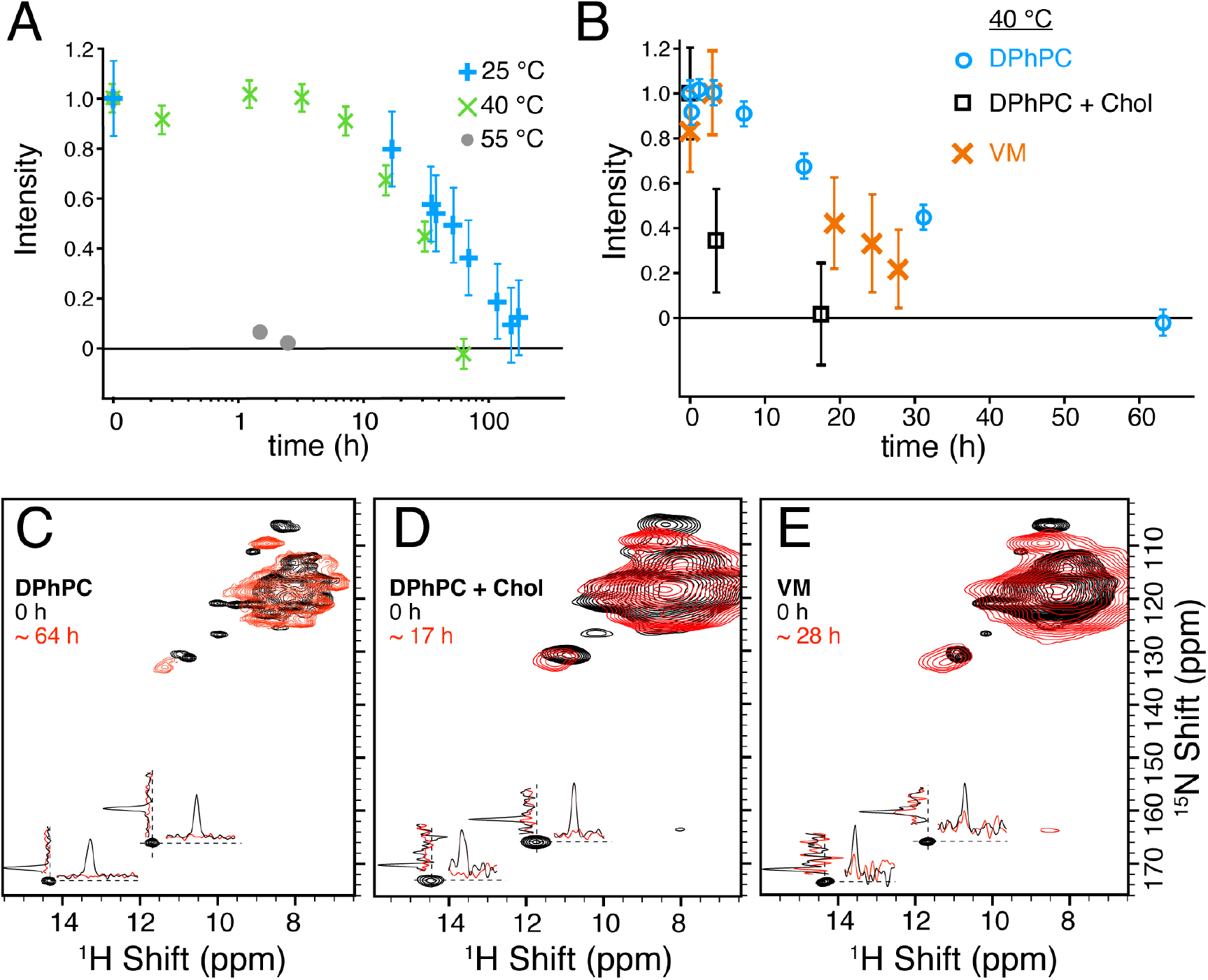
Kinetics of Rmt binding at different conditions. A) shows the disappearance of characteristic peaks (H37Nε2) of the apo (H)NH spectrum at three temperatures. The samples were initially prepared with 40 mM Rmt at 4 °C and then incubated by placing the rotor in a water bath at 25°C, 40 °C, or 55 °C. Two samples were used for the measurements. WT M2 was used for the 40 and 55 °C incubations, and H57Y M2 (a point mutation in the amphipathic domain) was used for the 25 °C measurements. Identical chemical shifts are observed for both samples. B) the average decay of the histidine side chain NHε2 signals measured at 40 °C is shown using DPhPC (circle), VM (cross) and DPhPC with 30% cholesterol (DPhPC+Chol, triangle). C)-E) shows the (H)NH spectra of M2 non-specific (black) and bound (red) for the three samples shown in B). These are the first (0h) and last points (timing as indicated in the caption) of the 40 °C kinetics for all the samples with the ^1^H and ^15^N slices shown for both of the H37 NHε2 peaks. Spectra in A) after 25 and 55 °C incubation were recorded at an 800 MHz Bruker Spectrometer with a 1.3 mm MAS probe spinning at 40 kHz MAS with VT gas set to 240 K. The spectra for the 40 °C kinetics using DPhPC were recorded on a 950 MHz Bruker spectrometer equipped with a 0.7 mm probe with VT gas set at 250 K and 80 kHz MAS. Spectra using VM and DPhPC with 30% cholesterol samples were recorded at 40 kHz MAS with a 1.3 mm rotor in a 800 MHz Bruker spectrometer setting VT gas at 240 K. In each case, the VT gas was set in order to reach a sample temperature during NMR data acquisition of about 278 K. The error bars shown only include errors from the spectrum noise and are shown as 2 times the root-mean-squared value.

An initial rate approximation was used to estimate the energy barrier (Ea) using the Arrhenius equation, 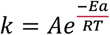 with k the rate, Ea the activation energy, R the gas constant and T the temperature. Pseudo first order kinetics is justified, considering that the high drug concentration remains relatively stable during the measurement. (A conservative estimate considering a protein concentration of ∼8.3 mM and ∼196 mM of lipids in the rotor and no concentration of the lipophilic drug in the membrane results in consumption of below 15% of the applied drug at saturation of all pore binding sites). Figure S6 shows the Arrhenius plot of ln(k) vs 1000/T were the slope gives direct access to the activation energy (slope = -Ea/R). Approximating the rates based on initial points until 50% decay of intensity, an activation energy of 122 ± 16 kJ per mol is found. The determination was repeated for individual protein cross-peaks, namely the side chain Nε2-Hε2 cross-peak of His37 and N-H cross-peak of Gly34 (Table S4-7). This is a relatively high activation energy, similar to the deactivation barrier encountered for enzymes,(43) and is therefore consistent with substantial change in the helical packing to accommodate drug entry into the pore. It suggests that the pore is not easily accessible to the drug in the apo state, consistent with previous reports indicating that the pore accessibility through the N-terminus is limited by residue V27, referred to as the secondary gate.(44)

Structural data suggests several processes in M2 that can account for this high energy barrier, namely structured water reorganization, imidazole-imidazole hydrogen bond disruption and pore opening. Studies on similar systems, where the pore contains a cluster of organized water molecules, have shown that the reorganization of water molecules necessitates ∼3-fold higher activation energies than bulk water,(45) but this is still only ∼30 kJ per mol.(46) X-ray structures suggest that during pore binding, Rmt displaces water molecules found at the N-terminus of the pore.(20, 21) In addition to the water displacement, Rmt pore binding disrupts the imidazole - imidazole hydrogen bond (27) yet this accounts for an energy of ∼10 kJ per mol.(47) Therefore helix repacking and transient opening of the N-terminus are likely to be responsible for the bulk of the activation energy.

Both the membrane composition and the M2 protein sequence can be expected to modulate the binding kinetics, in particular considering the differences in protein mobility and amantadine drug binding that were reported previously for TM and CD M2 constructs.(28) Recent molecular dynamics data provided additional evidence of the importance of the amphipathic helix for the pH gating mechanism.(48) We therefore reconstituted the CD construct into two additional membranes, one with DPhPC and 30% cholesterol and another in a mixture of lipids (see experimental section) referred to as the viral membrane (VM) composition, used previously by Hong and co-workers.(28) (H)NH and (H)CANH spectra of M2 reconstituted in DPhPC with 30% cholesterol have excellent ^1^H, ^15^N and ^13^C resolution similar to DPhPC membranes (Figure 4C-E, Figure S7 and Figure S8). Although broader lines are observed in the (H)NH and (H)CANH spectra recorded for the VM, the (H)NH spectra of all three samples show similar side chain histidine proton chemical shifts (HNε2) at 14.3 and 11.5 ppm (Figure 4E, Figure S7). These peaks are characteristic of the imidazole-imidazole hydrogen bond, which was detected in the DPhPC environment. Similar to DPhPC membranes, both, VM and DPhPC-Cholesterol at 4 °C show no pore binding upon addition of Rmt (Figure S9A-C, black). Pore binding at a 40 °C incubation temperature was tracked by measuring the intensity decay of the side chain histidine peaks from the (H)NH spectra using 40 kHz MAS (Figure 4B). The spectra from the beginning and end of the in incubation at 40 degrees is shown in Figure 4C-E) Interestingly, the DPhPC-Cholesterol sample showed faster pore binding kinetics compared to VM or DPhPC samples indicating that cholesterol as well as the lipid composition can modulate the binding kinetics, although in each case the binding occurred slowly over several hours. Cholesterol was reported to bind close to the peripheral binding site, near residues I39 to F47.(49, 50) The effect of cholesterol on binding kinetics is relatively small, but supports the consideration of the role of all membrane components.

In *in vitro* functional assays of M2 proton conduction, Rmt was observed to inhibit the protein after just 2 minutes of incubation. These functional assays are performed with a much higher LPR of about 200:1 by mass or about 4500 lipids per tetramer.(16) While this is in contrast to the slow kinetics observed in NMR samples, neither condition represents native membranes, in which M2 encounters a high density of other proteins such as hemagglutinin and neuraminidase in influenza particles, or a host of human proteins in endoplasmic reticulum, Golgi apparatus, or plasma membranes. The plasma membrane, from which M2 buds, is occupied by proteins at over 20 percent by area, and about 50% by mass.(51) It is therefore possible that such a high energy barrier is also encountered in a native context within host cellular membranes or viral particles.

However, it is also possible that the high energy barrier occurs only for the NMR preparations. In other words, the high protein and drug concentrations used in this study could potentially impact the conformational landscape of the protein. To begin to address this, we therefore recorded spectra using a high LPR of 20 (by mass) and recorded an apo spectrum. It displays the histidine peaks characteristic of imidazole-imidazole hydrogen bonding, with no significant changes in chemical shift detected (Figure S9A). We additionally found that the characteristic H37 peaks persist in different lipid compositions with an LPR of 20 by mass, corresponding to 1 M2 tetramer for 450 lipid molecules (Figure S9B-C). Unfortunately, the lower sensitivity of dilute samples prevented characterization of binding kinetics by NMR. For kinetics, low LPR of 1 to 1 by mass was used (25 to 1 by mole tetramer), which results in high NMR sensitivity, short acquisition time, and thus minimization of pore binding during acquisition. The high drug concentrations used in this study might also unduly influence the protein in the bound state in a non-specific manner. A 300 μM concentration of Rmt was therefore also applied, and confirmed to break the imidazole-imidazole hydrogen bonding interaction, and to shift the glycine amide peaks to the characteristic positions observed for pore-bound samples containing higher drug concentrations (Figure S9D).

Drug binding may occur *in vivo* at the lower pH of the Golgi or endosomes encountered by the virus during its life-cycle. The Rmt pore-binding kinetics are indeed faster at pH 6, which is close to the pH value of 6 to 6.7 pH found in Golgi (52). Upon addition of Rmt at pH 6, the histidine side chain peaks are almost fully decayed after 23 hours at 40 °C, which is 2-to 3-fold faster than at pH 7.8 (Figure S10). Previous studies have shown an interchain imidazole-imidazole or imidazole-imidazolium hydrogen bond at Histidine 37 (His37) position (27, 41). Based on M2 pKa measurements (30, 53-55), a pH of 6 results in population of multiple charge states at His37, including a significant population of the +3 charge state that is understood to be responsible for proton conduction. Loosening of the channel due to electrostatic repulsion in the positively charged His37 sidechains provides an explanation for the faster binding kinetics at low pH.

### Changes in the properties of M2 induced by pore binding prevent solubilization by DHPC detergent

Since we observed slow kinetics in lipid bilayer samples, we thought that this might be a limiting factor for drug binding in DHPC micelles as well. However, we could not prepare pore-bound samples in micelles by first binding Rmt in lipids, and then solubilizing in detergent. A solution of 300 mM 1,2-dihexanoyl-sn-glycerophosphocholine (DHPC) micelles was prepared and used to solubilize Rmt-bound M2 from the DPhPC lipid preparation (final detergent to lipid ratio of about 76). While some protein was solubilized, a large precipitate remained (Figure 5A). A second membrane sample of M2 without Rmt was fully solubilized in DHPC. Solution ^15^N-HSQC spectra were recorded for both supernatants (Figure 5B), and both matched the pore-apo M2 spectrum previously reported by Chou and co-workers.(24) Much lower sensitivity was observed in the supernatant from the bound sample, suggesting that only the remaining pore-apo protein could be solubilized. This was confirmed by MAS NMR measurements of the remaining pellet (Figure 5C), which resulted in a spectrum nearly identical to the rmt-bound spectrum in lipids. An explanation for this change in solubilization behavior could be that pore binding impacts the protein surface via repacking of the helices such that the solubility in DHPC micelles is decreased. Repacking of the helices due to drug binding in the pore has been suggested previously based on chemical shift perturbations that are observed far from the binding site.(26) The fact that a chimeric protein containing C-terminal residues from Influenza B M2 could be solubilized with drug in the pore, as reported by Chou and coworkers,(5) suggests that these effects may involve residues towards the C-terminus. Indeed, large chemical shift changes upon binding were reported for the chimeric protein for many residues, including those of the C-terminus, in particular at A40.(5)

**Figure 5.**
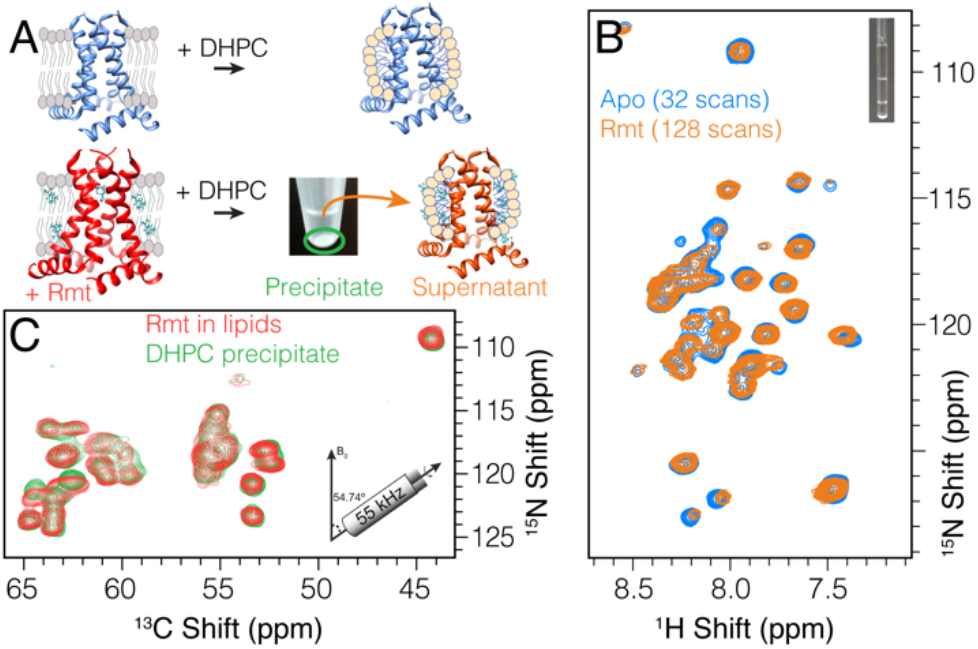
Pore binding results in channel restructuring. In A) a schematic view shows how the M2 tetramer was either dissolved directly with DHPC detergent (top, blue spectrum in B) or first bound with Rmt and then extracted with DHPC leaving a supernatant (orange spectrum in B) and an insoluble pellet (green spectrum in C). In B) the ^15^N-HSQC spectrum is shown for the soluble fraction after DHPC addition for the apo (blue) and pore bound (orange) samples. In C) the MAS spectrum of the pellet (green) is compared with the DPhPC lipid spectrum (red). Panel B) shows spectra recorded at a 600 MHz Bruker spectrometer using a cryoprobe. Panel C) shows projections of proton detected 3D (H)CANH spectra acquired at an 800 MHz spectrometer using 55 kHz MAS and a 1.3 mm rotor.

To our surprise, rather than finding a broad spectrum indicative of inhomogeneous aggregated protein, the pellet from the DHPC-detergent-treated bound sample showed nearly the same spectrum as the drug bound sample in lipid bilayers (Figure 5C). We could not detect deuterated lipids in the pellet (Figure S11). This suggests that bound tetramers likely pack close together in such a way that the sample remains stable in detergent, which would be consistent with the above explanation that M2 tetramer clustering might explain the observation of non-exponential drug binding kinetics.

## Conclusions

In summary, we show using proton detection in combination with ultra-fast MAS, that M2 can be kinetically trapped at low temperature in a pore-apo state for several different lipid membrane compositions. This allows measurement of non-specific CSP and tracking of binding kinetics with real time NMR under lipid bilayer conditions that resemble the viral membrane composition. Additionally, the formation of the imidazole-imidazole hydrogen bond was observed in each sample irrespective of the lipid composition. Nonetheless, we observed that the pore-binding kinetics are sensitive to the membrane composition and pH. The data suggest that drug binding causes a change in helix packing that impacts the aggregation properties as measured via dissolution of the membrane sample in DHPC micelles. This explains the lack of a pore-bound form in the resulting micellar solution. The high energy barrier to binding highlights the importance of kinetics when considering development of new inhibitors targeting the pore of M2, and underscores the potential role of kinetics more generally for pore-targeting small molecules.

## Acknowledgments

We thank James Chou for providing the expression plasmid, and Marcel Forster and Kai Xue for their feedback on the manuscript. We acknowledge financial support from the Max Planck Society and the DFG Emmy Noether program (grant AN 1316/1-1)

## Notes

### Competing Interest Statement

The authors have declared no competing interest.

