## Supplementary Information for "Real-time tracking of drug binding to Influenza A M2 reveals a high energy barrier"

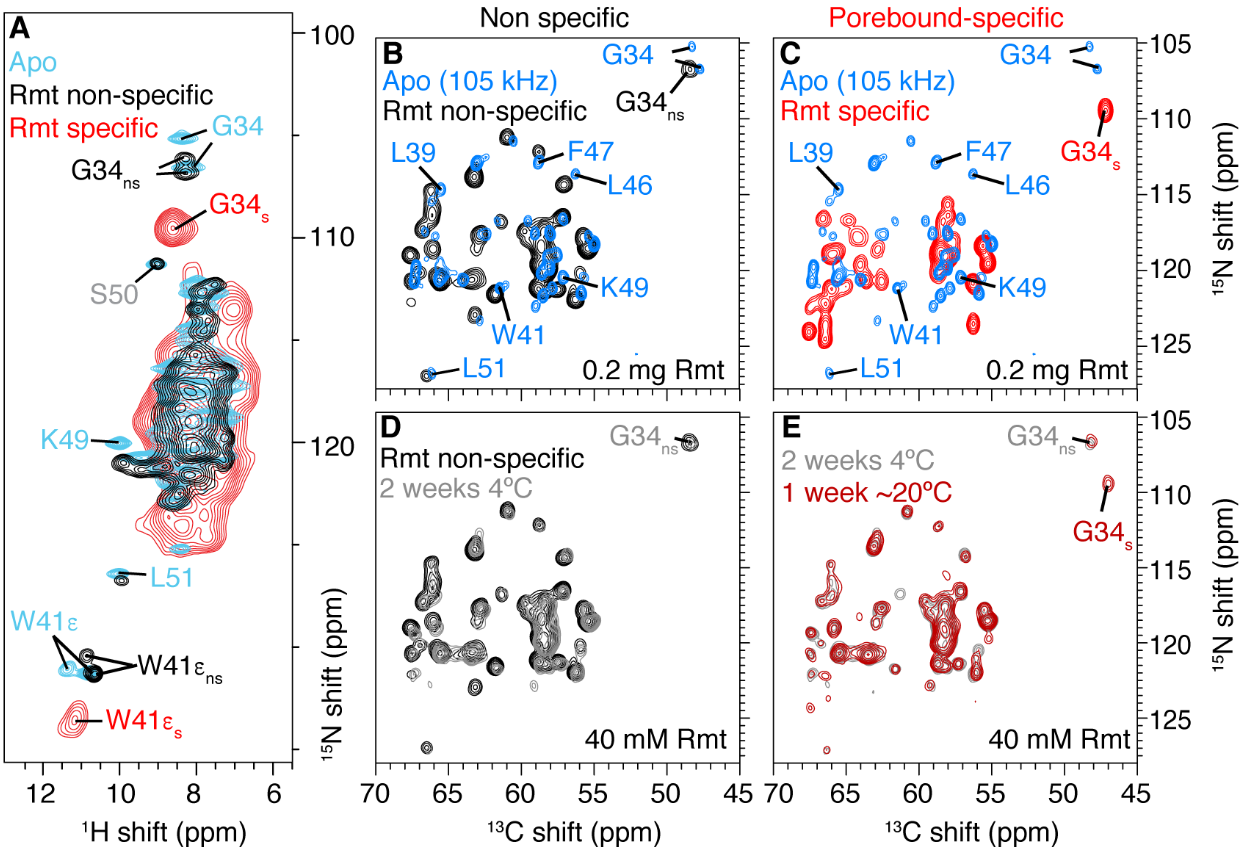

Figure S1: Tracking the pore binding process of M2 by MAS NMR. A) (H)NH spectra of apo M2 (cyan) with Rmt added demonstrating non-specific chemical shift perturbations (black) and specific chemical shift perturbations (red). B) to E) show CN projections of 3D (H)CANH spectra. In B) the apo spectrum (blue) is compared with non-specific chemical shift perturbations (black) by adding 0.2 mg of Rmt to the rotor (see methods for details). C) The same apo spectrum (blue) compared with the spectrum of the sample after the drug was bound to the pore, as indicated by the larger chemical shift changes (red). D) The non-specific spectrum of (B) compared with a sample incubated with 40 mM Rmt at 4 °C for 2 weeks (grey). E) The 2-week-incubated spectrum of (D) compared with a spectrum taken after an additional incubation for 1 week at 20 °C (dark red). Peaks characteristic of non-specific binding as well as peaks associated with specific binding are observed. Non-specific assignments are indicated with subscript “ns” in the spectra. The specific, pore bound, chemical shift assignments are indicated with subscript “s”. Unless indicated, spectra were recorded at 55 kHz on an 850 MHz wide-bore spectrometer using a 4-channel 1.3 mm MAS probe and VT gas at 240 K (panel A, cyan and black; panel B, black) or on an 800 MHz wide-bore spectrometer using a 3-channel 1.3 mm MAS probe and VT gas at 235 K (panel A, red; panels C-E). The 105 kHz spectrum (panel B-C, blue) was recorded on a 950 MHz spectrometer with VT gas set to 260 K. In each the estimated sample temperature is approximately 15 - 20 °C.

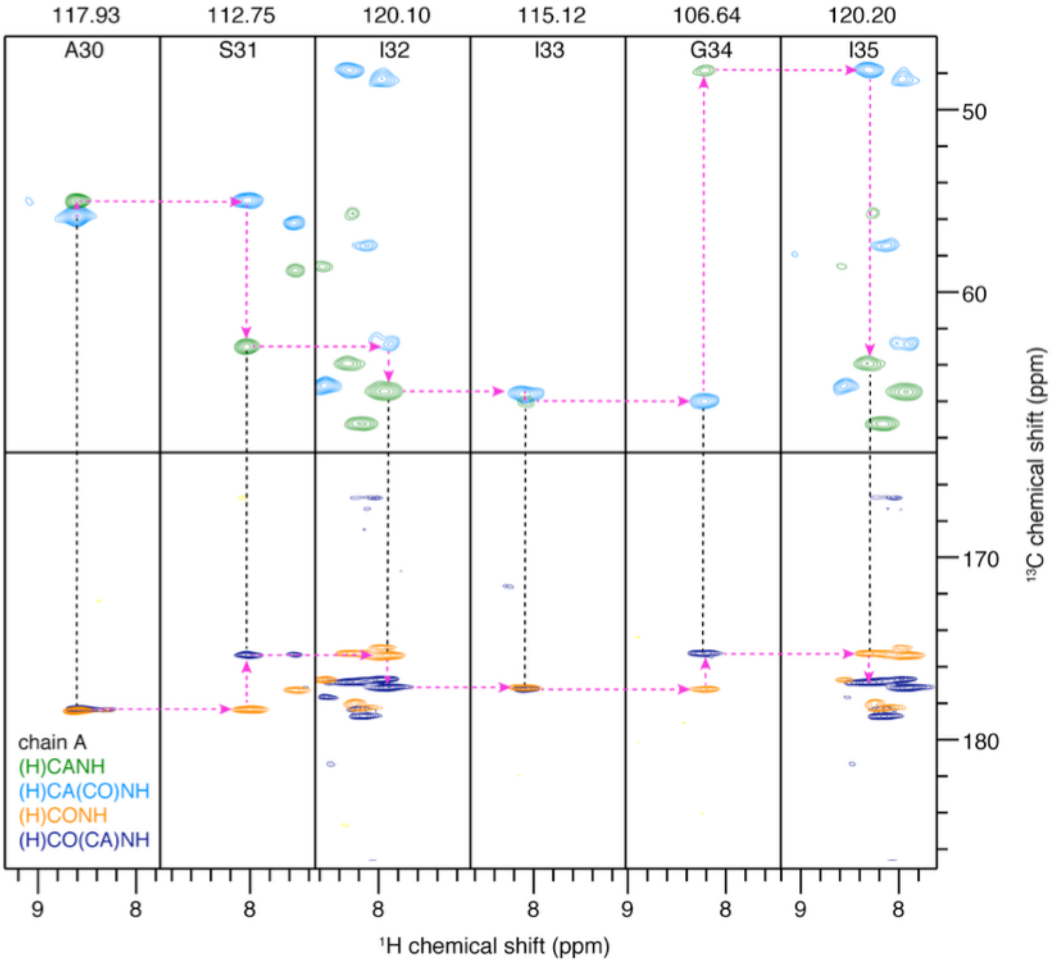

Figure S2: M2 chain A residue-specific assignment. Strip plots showing the connection from residue A30 to I35 in two pairs of spectra, linking through either CA or CO, with detection at amide protons.(1)

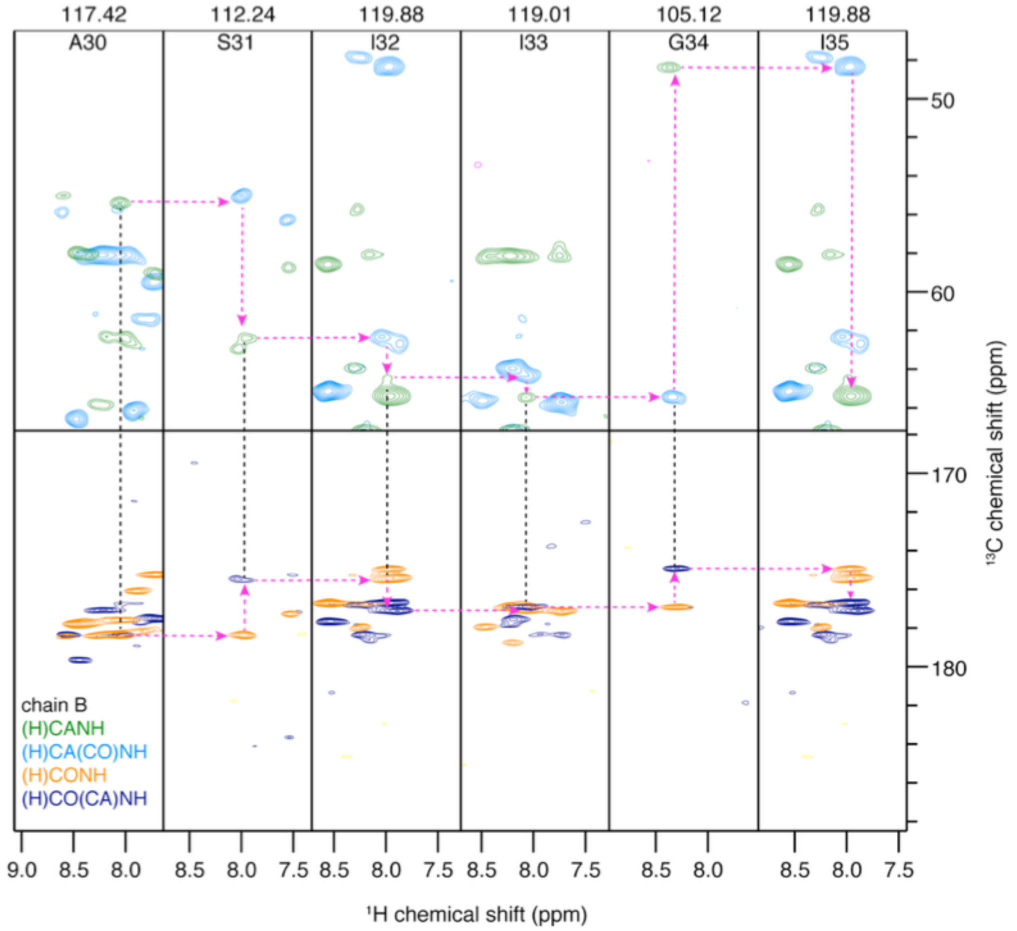

Figure S3: M2 chain B residue-specific assignment. Strip plots showing the connection from residues A30 to I35 in two pairs of spectra, linking through either CA or CO, with detection at amide protons. [1]

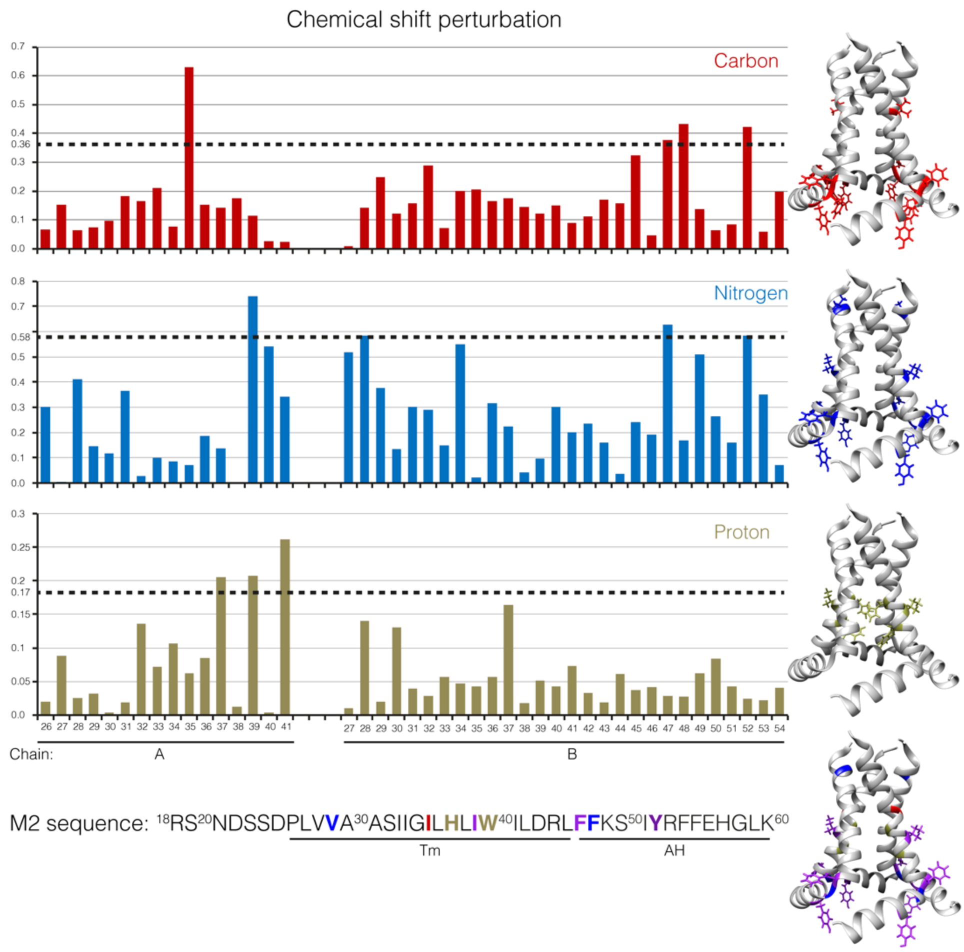

Figure S4: Individual chemical shift perturbation (CSP) observed for Rmt non-specific binding in M2. Carbon, nitrogen, and proton CSP are shown in red, blue, and gold, respectively. The amino acid sequence of M2 is shown at the bottom, and the structures at the right (based on pdb 2L0J) are colored for residues with CSP above 2 times the root mean squared CSP, as indicated by the dotted lines.

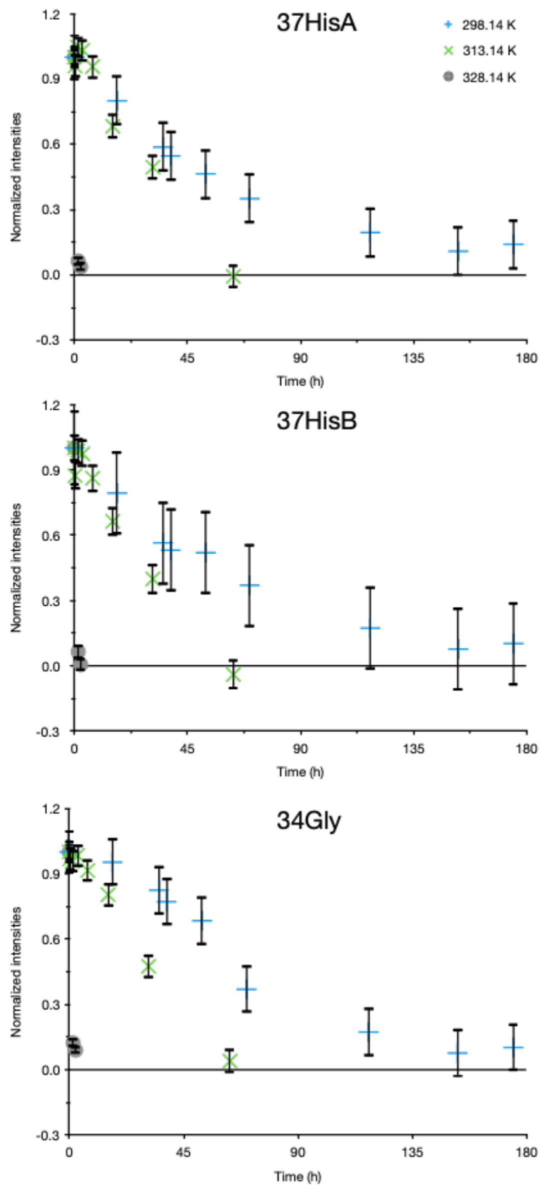

Figure S5: Time dependence of rimantadine binding at 25 °C (+), 40 °C(x) and 55 °C (circles) tracked by real-time NMR. The different panels show the binding kinetics for different residues (H37A-HNε2, H37B-HNε2 and G34). Two distinct samples were used, WT M2 for the 40 and 55 °C measurement and the H57Y mutant for the 25 °C kinetics.

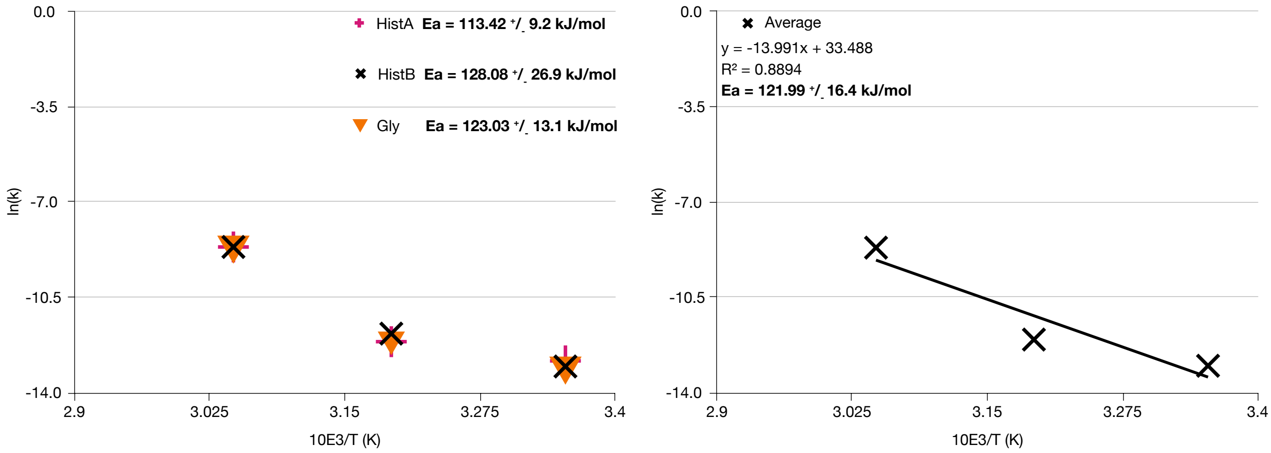

Figure S6: Activation energy required for Rmt pore-binding in M2 in DPhPC membranes. The Left panel shows the Arrhenius plot and the activation energy as determined from peak intensities of different residues (see table S4, S5 and S6 for the intensities): G34 (triangle), H37A-HNε2 (+) and H37B-HNε2 (x). The right panel shows the Arrhenius plot for the average of the three residues, Gly, HistA and HistB. WT M2 was used for data at 40 and 55 °C and the H57Y mutant was used at 20 °C. All samples were initially incubated at 5°C in the presence of 40mM Rmt.

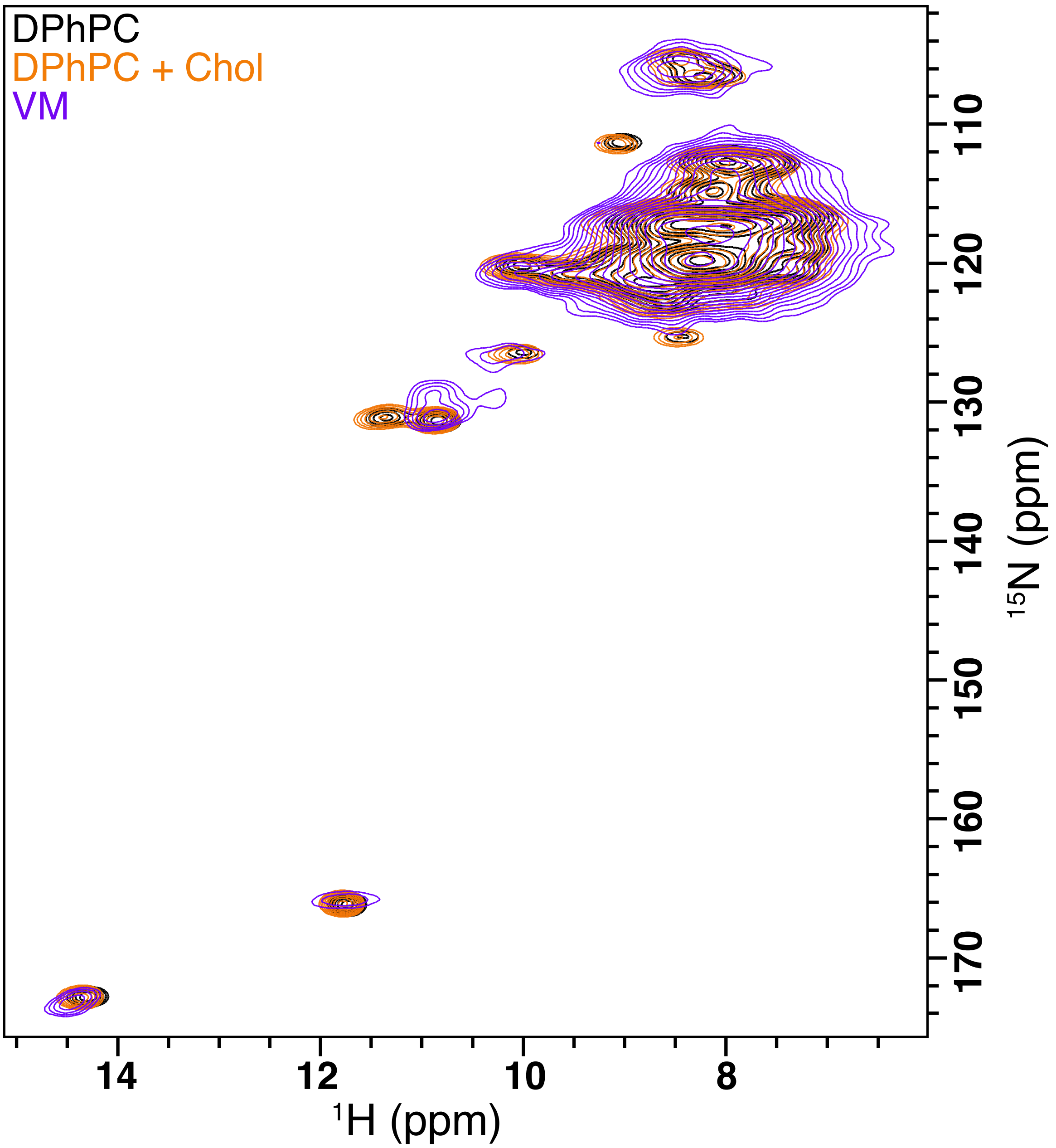

Figure S7: The histidine side chain shows that M2 adopts a dimer of dimer conformation in different lipids membranes. Comparison of the M2 conductance domain construct reconstituted in DPhPC (black), DPhPC+cholesterol (orange) and VM (purple). The presence of both H37A-HNε2 ($^{1}$H: ~11.8 ppm, $^{15}$N: ~167 ppm) and H37B-HNε2 ($^{1}$H: ~14.2 ppm, $^{15}$N: ~174 ppm) indicates the presence of a dimer of dimers as we reported previously.(2) This is further supported by the 2 side chain tryptophan peaks observed at ~130 ppm nitrogen and 11 ppm proton since there is only 1 tryptophan, W41, in the amino acid sequence of M2. Glycine peaks are broader in the VM membranes. This might be due to inhomogeneity of the sample and/or dynamics. The spectra were recorded at 55 kHz MAS on a 18.8 Tesla Bruker spectrometer using a three channel probe and with the cooling gas temperature set to 235 K, which corresponds to a sample temperature of about 293 K (20 °C). All the (H)NH spectra where processed using 8 ms in the direct dimension ($^{1}$H), and 10 ms in the indirect dimension ($^{15}$N). We used a cosine squared window function in both dimensions during processing.

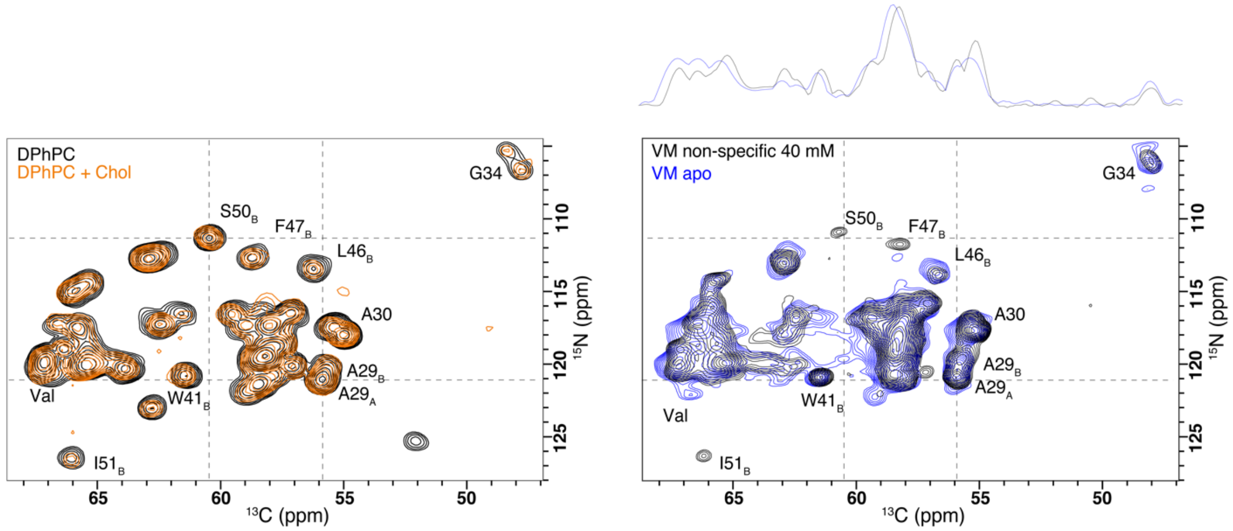

Figure S8: Comparison of carbon and nitrogen 2D projections from 3D (H)CANH spectra of the M2 conductance domain construct reconstituted in DPhPC (left, black), DPhPC with 30% cholesterol (DPhPC+Chol in orange) and VM (blue). (right) Non-specific changes after addition of 40 mM rimantadine are compared in black. No changes are observed that would indicate a change in secondary structure of the M2 protein. Note the consistency of peak doubling in the Gly and Ala peaks, and consistent peak positions for C-terminal residues including residues of the amphipathic helix (L46-I51). Several peaks are missing (undetected) or weak in the VM sample. It is not clear if this arises due to sample heterogeneity in the more complex lipids, or from changes in dynamics. The spectra were recorded at 55 kHz MAS on an 18.8 Tesla Bruker spectrometer using a three-channel probe and with cooling gas set temperature of 235 K, which corresponds to a sample temperature of about 293 K (20 °C).The spectra were processed using 8 ms on $^{1}$H, 6 ms on $^{13}$C and 10 ms on $^{15}$N. A cosine squared window function was applied in all dimensions during processing.

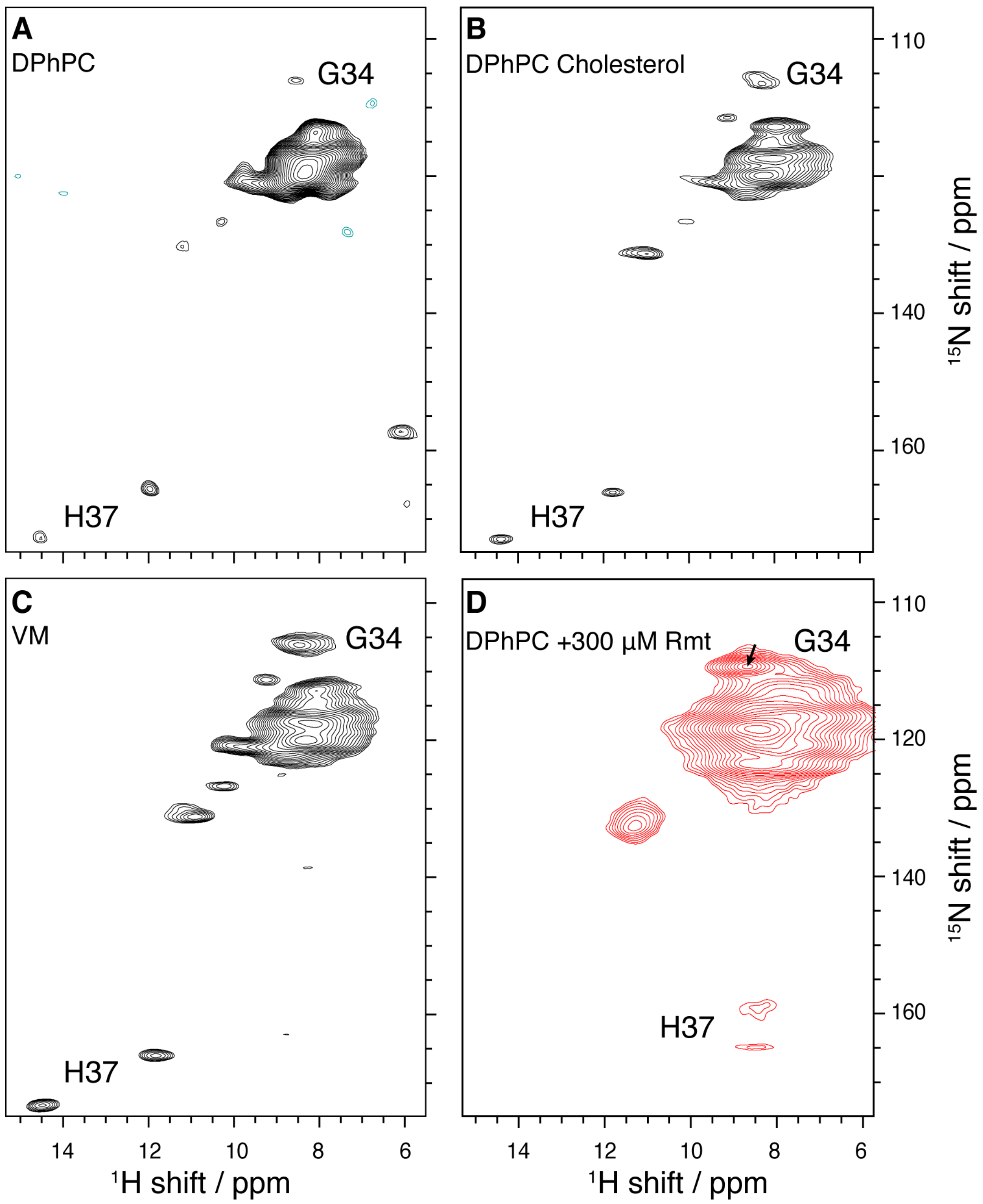

Figure S9: M2 retains similar fold when reconstituted using high lipid protein ratios, and drug binding results in similar changes to the spectrum with a lower, 300 µM concentration. Panels A to C show the (H)NH spectra of M2 reconstituted in DPhPC membranes (A), DPhPC with 30% cholesterol (B) and VM membranes (C) each using a 20 to 1 lipid to protein ratio by mass, which is about 460 lipids per tetramer . Panel (D) shows a spectrum of M2 treated with 300 µM Rmt rather than the usual 40 mM concentration. The sample was incubated at 55 °C for about 5 hours and adopts the characteristic pore-bound spectrum without the His-His hydrogen bond, and with a clear change in the chemical shift of G34, as indicated by the black arrow. For the incubation, sufficient drug was ensured by use of a 200 µL buffer volume. The spectra were acquired on an 800 MHz Bruker spectrometer and sampled to 21 ms in the direct dimension ($^{1}$H) and 10 ms on the indirect dimension ($^{15}$N) corresponding roughly to a total of 2 to 4 days per spectrum. All the spectra were processed using 5 ms in the direct dimension ($^{1}$H) and 10 ms in the indirect dimension ($^{15}$N) applying a cosine squared apodization function in both dimensions. Baseline correction using qfil was applied in the direct dimension ($^{1}$H).

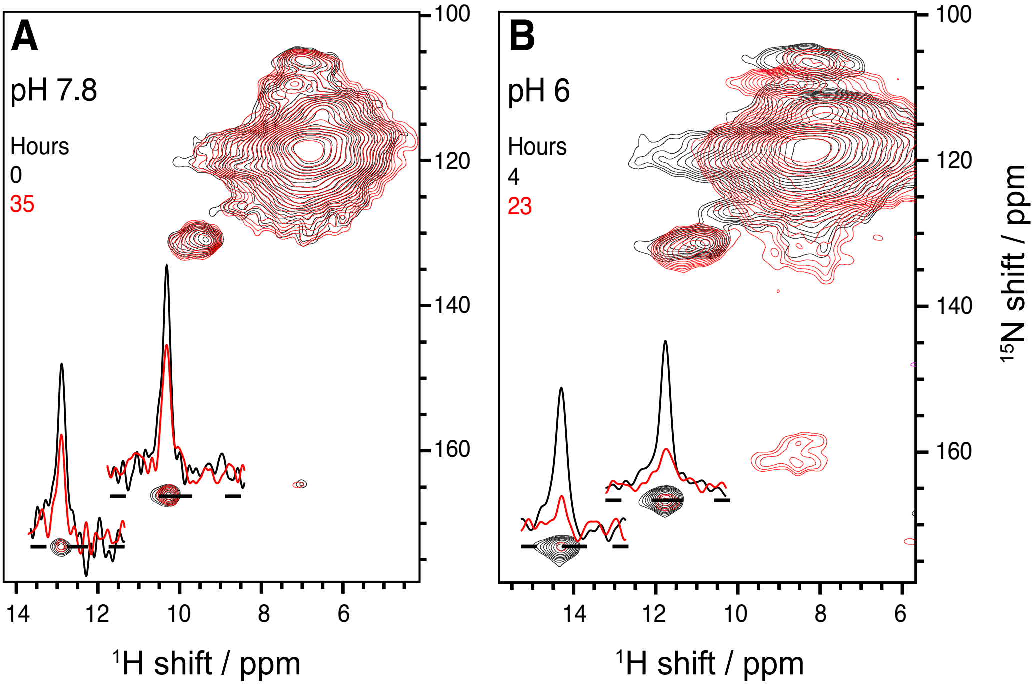

Figure S10: Rmt pore binding kinetics at different pH followed by real time NMR. Panel A) shows the (H)NH spectra of M2 incubated with 40 mM Rmt at pH 7.8 at 4 °C (black) and after incubation at 25 °C for ~35 hours (red). Panel B) shows the (H)NH spectra of M2 with 40 mM Rmt at pH 6 (similar to the pH found in the Golgi compartment) after 4 hours incubation at 25 °C (black) and 23 hours (red). The spectra in A were recorded on an 800 MHz at Bruker spectrometer equipped with a 1.3 mm probe spinning at 40 kHz and VT set at 240 K. Both spectra displayed in B were recorded using a 600 MHz Bruker spectrometer equipped with a 1.3 mm probe. The VT gas was set at 240 K and the MAS frequency at 40 kHz. All the spectra were processed using 8 ms in the direct dimension ($^1$H) and 8 ms in the indirect dimension ($^{15}$N).

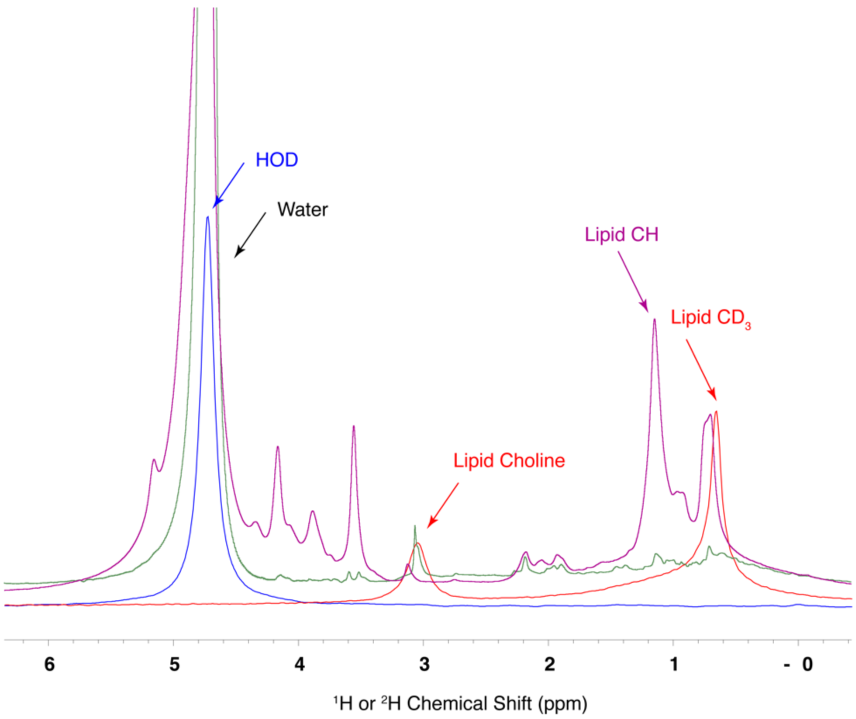

Figure S11: Single pulse proton and deuterium spectra of M2 samples in deuterated DPHPC lipids (purple and red, respectively) or after treatment with DHPC detergent (green and blue, respectively). While both choline and aliphatic signals are clearly observed in the deuterium spectrum of the lipid bilayer preparation, only HOD was detected in the pellet that remained after treatment with DHPC detergent. Since both DPHPC and DHPC have similar chemical groups, comparison of the proton spectra does not readily distinguish between lipid and detergent. Data were acquired at a 600 MHz spectrometer.

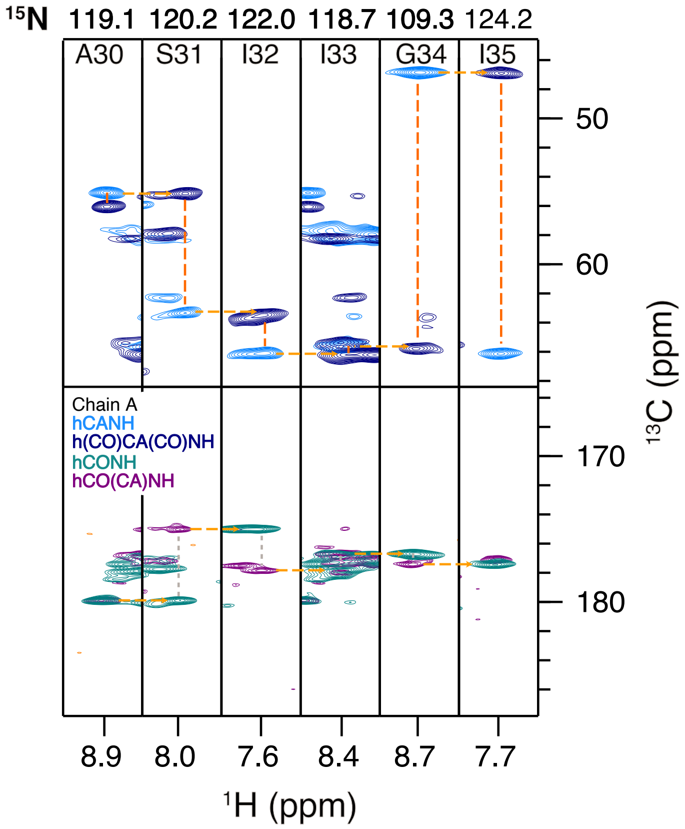

Figure S12: Rmt pore bound M2 chain A residue-specific assignment. Strip plots showing the connection from residues A30 to I35 in two pairs of spectra, linking through either CA or CO, with detection at amide protons. [1]

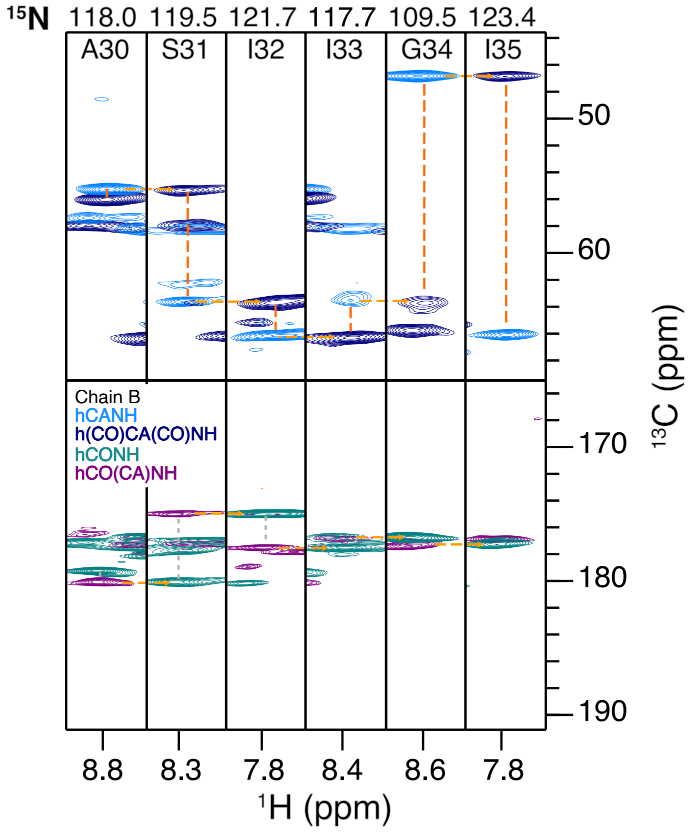

Figure S13: Rmt pore bound M2 chain B residue-specific assignment. Strip plots showing the connection from residues A30 to I35 in two pairs of spectra, linking through either CA or CO, with detection at amide protons. [1]

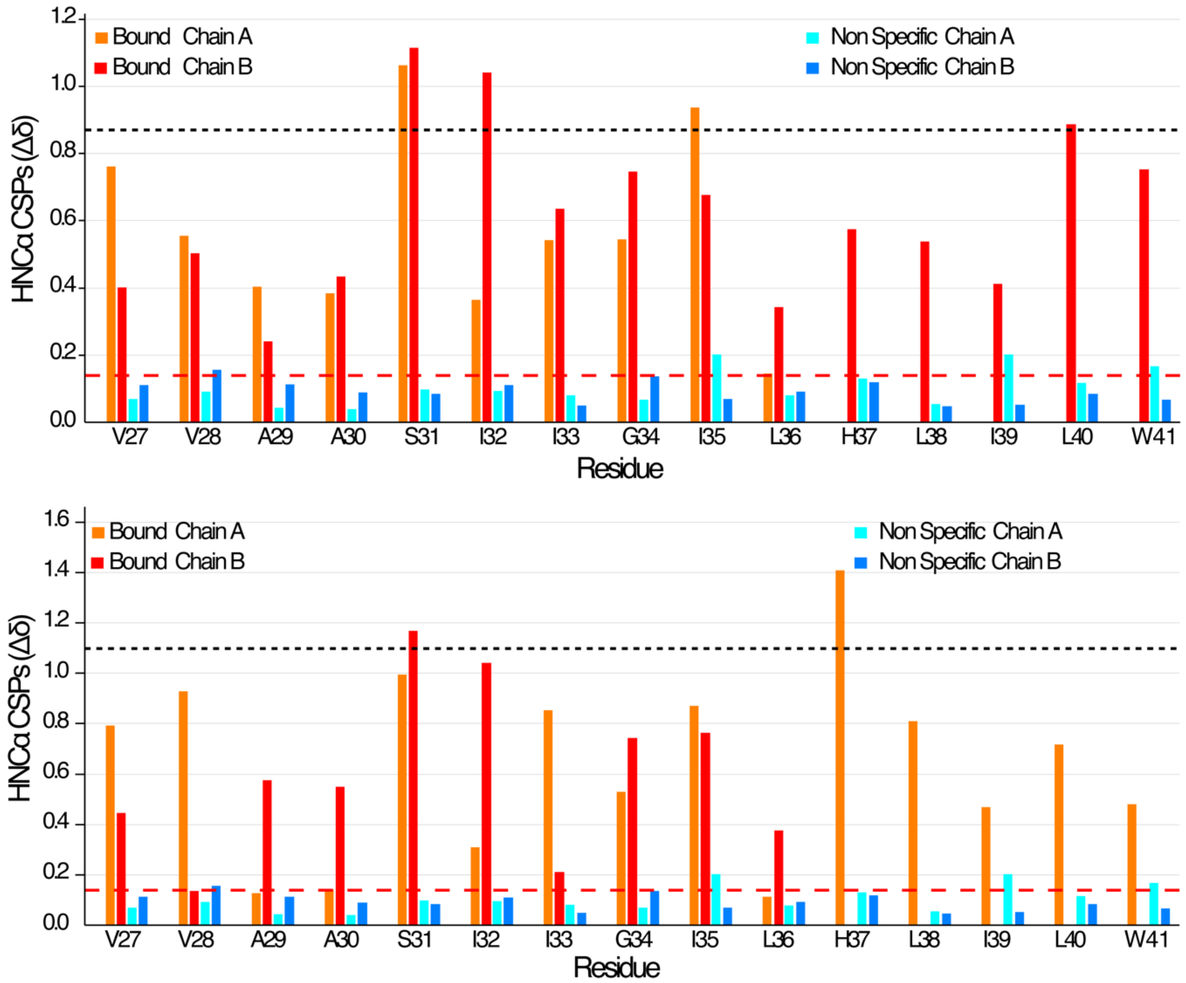

Figure S14: Comparison of chemical shift perturbation (CSP) observed for specific and non-specific binding in M2. (top) ΔδCSP are shown in orange and red for specific changes in chain A and B, respectively. ΔδCSP are shown in cyan and blue for non-specific changes in chain A and B, respectively. Dashed black and red lines indicate 3 times the root mean square deviation of bound vs apo and non-specific vs apo, respectively. (bottom) Same as the top plot, but with the definition of chain A and B for specific pore bound shifts interchanged.

Table S1. Assigned chemical shifts for apo $M2_{18-60}$.

| Chain A | | | | Chain B | | | |
|---|---|---|---|---|---|---|---|
| residue number | Amino acid | Atom | chemical shift | residue number | Amino acid | Atom | chemical shift |
| 25 | P | C | 177.27 | 25 | P | C | * |
| 25 | P | CA | 65.70 | 25 | P | CA | * |
| 26 | L | C | 178.28 | 26 | L | C | 178.26 |
| 26 | L | CA | 57.53 | 26 | L | CA | * |
| 26 | L | CB | 40.32 | 26 | L | CB | * |
| 26 | L | HN | 7.71 | 26 | L | HN | * |
| 26 | L | N | 118.65 | 26 | L | N | * |
| 27 | V | C | 178.75 | 27 | V | C | 178.57 |
| 27 | V | CA | 67.24 | 27 | V | CA | 66.43 |
| 27 | V | CB | 31.42 | 27 | V | CB | * |
| 27 | V | HN | 8.16 | 27 | V | HN | 7.27 |
| 27 | V | N | 120.33 | 27 | V | N | 118.83 |
| 28 | V | C | 177.83 | 28 | V | C | 177.98 |
| 28 | V | CA | 67.12 | 28 | V | CA | 66.77 |
| 28 | V | CB | 31.39 | 28 | V | CB | * |
| 28 | V | HN | 8.22 | 28 | V | HN | 7.88 |
| 28 | V | N | 119.43 | 28 | V | N | 120.71 |
| 29 | A | C | 178.41 | 29 | A | C | 178.33 |
| 29 | A | CA | 55.90 | 29 | A | CA | 55.72 |
| 29 | A | CB | 18.60 | 29 | A | CB | * |
| 29 | A | HN | 8.40 | 29 | A | HN | 8.25 |
| 29 | A | N | 121.12 | 29 | A | N | 120.09 |
| 30 | A | C | 178.34 | 30 | A | C | 178.39 |
| 30 | A | CA | 55.00 | 30 | A | CA | 55.49 |
| 30 | A | CB | 18.63 | 30 | A | CB | 18.39 |
| 30 | A | HN | 8.60 | 30 | A | HN | 8.06 |
| 30 | A | N | 117.92 | 30 | A | N | 117.40 |
| 31 | S | C | 175.38 | 31 | S | C | 175.41 |
| 31 | S | CA | 62.89 | 31 | S | CA | 62.38 |
| 31 | S | CB | 63.44 | 31 | S | CB | * |
| 31 | S | HN | 8.01 | 31 | S | HN | 7.96 |
| 31 | S | N | 112.70 | 31 | S | N | 112.17 |
| 32 | I | C | 177.14 | 32 | I | C | 177.06 |
| 32 | I | CA | 65.53 | 32 | I | CA | 62.86 |
| 32 | I | CB | 37.81 | 32 | I | CB | 37.39 |
| 32 | I | HN | 7.93 | 32 | I | HN | 7.96 |
| 32 | I | N | 120.14 | 32 | I | N | 119.87 |
| 33 | I | C | 177.23 | 33 | I | C | 176.92 |
| 33 | I | CA | 66.03 | 33 | I | CA | 65.44 |
| 33 | I | CB | 37.17 | 33 | I | CB | 37.22 |

| | | | | | | | |
|---|---|---|---|---|---|---|---|
| 33 | I | HN | 8.09 | 33 | I | HN | 8.06 |
| 33 | I | N | 115.10 | 33 | I | N | 119.00 |
| 34 | G | C | 175.27 | 34 | G | C | 174.94 |
| 34 | G | CA | 47.80 | 34 | G | CA | 48.26 |
| 34 | G | HN | 8.19 | 34 | G | HN | 8.31 |
| 34 | G | N | 106.62 | 34 | G | N | 105.18 |
| 35 | I | C | 176.86 | 35 | I | C | 176.71 |
| 35 | I | CA | 63.88 | 35 | I | CA | 64.63 |
| 35 | I | CB | 37.35 | 35 | I | CB | 37.60 |
| 35 | I | HN | 8.32 | 35 | I | HN | 7.97 |
| 35 | I | N | 120.26 | 35 | I | N | 119.93 |
| 36 | L | C | 177.58 | 36 | L | C | 177.66 |
| 36 | L | CA | 58.12 | 36 | L | CA | 58.56 |
| 36 | L | CB | 41.73 | 36 | L | CB | 41.82 |
| 36 | L | HN | 8.20 | 36 | L | HN | 8.54 |
| 36 | L | N | 119.37 | 36 | L | N | 119.81 |
| 37 | H | C | 176.74 | 37 | H | C | 175.23 |
| 37 | H | CA | 62.33 | 37 | H | CA | 59.48 |
| 37 | H | CB | 32.14 | 37 | H | CB | 31.10 |
| 37 | H | HN | 8.00 | 37 | H | HN | 8.37 |
| 37 | H | N | 117.30 | 37 | H | N | 116.50 |
| 38 | L | C | 176.74 | 38 | L | C | 177.51 |
| 38 | L | CA | 58.11 | 38 | L | CA | 59.00 |
| 38 | L | CB | 40.59 | 38 | L | CB | 42.40 |
| 38 | L | HN | 8.10 | 38 | L | HN | 7.71 |
| 38 | L | N | 119.58 | 38 | L | N | 117.27 |
| 39 | I | C | 177.08 | 39 | I | C | 177.02 |
| 39 | I | CA | 65.81 | 39 | I | CA | 65.41 |
| 39 | I | CB | 37.28 | 39 | I | CB | 37.79 |
| 39 | I | HN | 8.26 | 39 | I | HN | 8.02 |
| 39 | I | N | 117.41 | 39 | I | N | 114.43 |
| 40 | L | C | * | 40 | L | C | 178.80 |
| 40 | L | CA | 58.15 | 40 | L | CA | 58.50 |
| 40 | L | CB | 40.53 | 40 | L | CB | 41.35 |
| 40 | L | HN | 7.73 | 40 | L | HN | 8.82 |
| 40 | L | N | 118.84 | 40 | L | N | 121.28 |
| 41 | W | C | 178.54 | 41 | W | C | 178.16 |
| 41 | W | CA | 62.80 | 41 | W | CA | 61.41 |
| 41 | W | CB | 28.09 | 41 | W | CB | 27.38 |
| 41 | W | HN | 8.51 | 41 | W | HN | 9.46 |
| 41 | W | N | 123.00 | 41 | W | N | 120.81 |
| 42 | I | C | 177.95 | 42 | I | C | 177.77 |
| 42 | I | CA | 65.64 | 42 | I | CA | 66.58 |
| 42 | I | CB | 37.45 | 42 | I | CB | 37.75 |

| | | | | | | | |
|---|---|---|---|---|---|---|---|
| 42 | I | HN | 8.81 | 42 | I | HN | 7.84 |
| 42 | I | N | 119.54 | 42 | I | N | 117.14 |
| 43 | L | C | * | 43 | L | C | 179.65 |
| 43 | L | CA | 58.21 | 43 | L | CA | 57.98 |
| 43 | L | CB | * | 43 | L | CB | 41.72 |
| 43 | L | HN | 8.49 | 43 | L | HN | 8.45 |
| 43 | L | N | 119.15 | 43 | L | N | 117.25 |
| | | | | 44 | D | C | 178.45 |
| | | | | 44 | D | CA | 57.78 |
| | | | | 44 | D | CB | 42.62 |
| | | | | 44 | D | HN | 9.06 |
| | | | | 44 | D | N | 120.83 |
| | | | | 45 | R | C | 178.30 |
| | | | | 45 | R | CA | 57.05 |
| | | | | 45 | R | CB | 30.52 |
| | | | | 45 | R | HN | 8.54 |
| | | | | 45 | R | N | 116.32 |
| | | | | 46 | L | C | 177.26 |
| | | | | 46 | L | CA | 56.27 |
| | | | | 46 | L | CB | 42.14 |
| | | | | 46 | L | HN | 7.80 |
| | | | | 46 | L | N | 113.32 |
| | | | | 47 | F | C | 175.34 |
| | | | | 47 | F | CA | 58.76 |
| | | | | 47 | F | HN | 7.54 |
| | | | | 47 | F | N | 112.64 |
| | | | | 48 | F | C | 175.60 |
| | | | | 48 | F | CA | 61.52 |
| | | | | 48 | F | CB | 39.51 |
| | | | | 48 | F | HN | 7.37 |
| | | | | 48 | F | N | 116.45 |
| | | | | 49 | K | C | 175.65 |
| | | | | 49 | K | CA | 57.14 |
| | | | | 49 | K | CB | 30.30 |
| | | | | 49 | K | HN | 9.96 |
| | | | | 49 | K | N | 120.11 |
| | | | | 50 | S | C | 174.68 |
| | | | | 50 | S | CA | 60.47 |
| | | | | 50 | S | CB | 66.25 |
| | | | | 50 | S | HN | 9.02 |
| | | | | 50 | S | N | 111.29 |
| | | | | 51 | I | C | 176.09 |
| | | | | 51 | I | CA | 66.10 |
| | | | | 51 | I | CB | 38.12 |

| | | | | | | | |
|---|---|---|---|---|---|---|---|
| | | | | 51 | I | HN | 9.93 |
| | | | | 51 | I | N | 126.46 |
| | | | | 52 | Y | C | 178.93 |
| | | | | 52 | Y | CA | 62.74 |
| | | | | 52 | Y | HN | 7.91 |
| | | | | 52 | Y | N | 117.34 |
| | | | | 53 | R | CA | 58.90 |
| | | | | 53 | R | CB | 29.49 |
| | | | | 53 | R | HN | 8.54 |
| | | | | 53 | R | N | 121.96 |
| | | | | 54 | F | CA | 61.02 |
| | | | | 54 | F | HN | 7.31 |
| | | | | 54 | F | N | 120.64 |

Table S2. Assigned chemical shifts for non-specific binding of Rmt to $M2_{18-60}$ using 40 mM Rmt.

| Chain A | | | | Chain B | | | |
|---|---|---|---|---|---|---|---|
| Residue number | Residue | Atom | Chemical shift | Residue number | Residue | Atom | Chemical shift |
| 25 | P | C | 177.70 | 25 | P | C | * |
| 26 | L | C | 178.13 | 26 | L | C | 178.31 |
| 26 | L | CA | 57.54 | 26 | L | CA | * |
| 26 | L | H | 7.75 | 26 | L | H | * |
| 26 | L | N | 118.91 | 26 | L | N | * |
| 27 | V | C | 178.51 | 27 | V | C | 178.79 |
| 27 | V | CA | 67.10 | 27 | V | CA | 66.42 |
| 27 | V | H | 8.23 | 27 | V | H | 7.30 |
| 27 | V | N | 120.35 | 27 | V | N | 119.44 |
| 28 | V | C | 178.16 | 28 | V | C | 177.77 |
| 28 | V | CA | 67.07 | 28 | V | CA | 66.91 |
| 28 | V | H | 8.22 | 28 | V | H | 8.02 |
| 28 | V | N | 119.00 | 28 | V | N | 121.22 |
| 29 | A | C | 178.40 | 29 | A | C | 178.36 |
| 29 | A | CA | 55.83 | 29 | A | CA | 55.44 |
| 29 | A | H | 8.35 | 29 | A | H | 8.24 |
| 29 | A | N | 121.29 | 29 | A | N | 119.75 |
| 30 | A | C | 178.36 | 30 | A | C | 178.38 |
| 30 | A | CA | 54.92 | 30 | A | CA | 55.27 |
| 30 | A | H | 8.60 | 30 | A | H | 8.18 |
| 30 | A | N | 118.06 | 30 | A | N | 117.32 |
| 31 | S | C | 175.26 | 31 | S | C | * |
| 31 | S | CA | 62.83 | 31 | S | CA | 62.52 |
| 31 | S | H | 8.06 | 31 | S | H | 7.99 |
| 31 | S | N | 113.09 | 31 | S | N | 112.45 |
| 32 | I | C | 177.08 | 32 | I | C | 177.10 |
| 32 | I | CA | 65.33 | 32 | I | CA | 64.22 |
| 32 | I | H | 7.79 | 32 | I | H | 8.01 |
| 32 | I | N | 120.22 | 32 | I | N | 120.14 |
| 33 | I | C | 177.24 | 33 | I | C | 176.88 |
| 33 | I | CA | 65.83 | 33 | I | CA | 65.39 |
| 33 | I | H | 8.13 | 33 | I | H | 8.13 |
| 33 | I | N | 115.07 | 33 | I | N | 119.25 |
| 34 | G | C | 175.25 | 34 | G | C | 174.89 |
| 34 | G | CA | 47.78 | 34 | G | CA | 48.18 |
| 34 | G | H | 8.31 | 34 | G | H | 8.40 |
| 34 | G | N | 106.52 | 34 | G | N | 105.70 |
| 35 | I | C | 177.06 | 35 | I | C | 176.62 |
| 35 | I | CA | 63.29 | 35 | I | CA | 65.22 |

| | | | | | | | |
|---|---|---|---|---|---|---|---|
| 35 | I | H | 8.26 | 35 | I | H | 7.92 |
| 35 | I | N | 120.21 | 35 | I | N | 120.03 |
| 36 | L | C | 177.47 | 36 | L | C | * |
| 36 | L | CA | 57.98 | 36 | L | CA | 58.40 |
| 36 | L | H | 8.27 | 36 | L | H | 8.49 |
| 36 | L | N | 119.59 | 36 | L | N | 119.47 |
| 37 | H | C | 176.65 | 37 | H | C | 175.20 |
| 37 | H | CA | 62.24 | 37 | H | CA | 59.36 |
| 37 | H | H | 8.17 | 37 | H | H | 8.53 |
| 37 | H | HE2 | 11.67 | 37 | H | HE2 | 14.34 |
| 37 | H | N | 117.16 | 37 | H | N | 116.67 |
| 37 | H | NE2 | 165.90 | 37 | H | NE2 | 173.11 |
| 38 | L | C | * | 38 | L | C | 177.53 |
| 38 | L | CA | 57.97 | 38 | L | CA | 58.91 |
| 38 | L | H | 8.06 | 38 | L | H | 7.73 |
| 38 | L | N | 119.53 | 38 | L | N | 117.31 |
| 39 | I | C | 177.04 | 39 | I | C | 177.03 |
| 39 | I | CA | 65.70 | 39 | I | CA | 65.55 |
| 39 | I | H | 8.07 | 39 | I | H | 7.97 |
| 39 | I | N | 116.67 | 39 | I | N | 114.31 |
| 40 | L | C | 177.99 | 40 | L | C | 178.72 |
| 40 | L | CA | 58.08 | 40 | L | CA | 58.36 |
| 40 | L | H | 7.73 | 40 | L | H | 8.88 |
| 40 | L | N | 118.27 | 40 | L | N | 120.99 |
| 41 | W | CA | 62.77 | 41 | W | C | 178.11 |
| 41 | W | H | 8.72 | 41 | W | CA | 61.34 |
| 41 | W | HE1 | 11.06 | 41 | W | H | 9.53 |
| 41 | W | N | 122.60 | 41 | W | HE1 | 10.74 |
| 41 | W | NE1 | 130.33 | 41 | W | N | 121.00 |
| | | | | 41 | W | NE1 | 131.10 |
| | | | | 42 | I | C | 177.75 |
| | | | | 42 | I | CA | 66.42 |
| | | | | 42 | I | H | 7.82 |
| | | | | 42 | I | N | 116.92 |
| | | | | 43 | L | C | 179.57 |
| | | | | 43 | L | CA | 57.85 |
| | | | | 43 | L | H | 8.46 |
| | | | | 43 | L | N | 117.03 |
| | | | | 44 | D | C | 178.53 |
| | | | | 44 | D | CA | 57.71 |
| | | | | 44 | D | H | 9.13 |
| | | | | 44 | D | N | 120.85 |
| | | | | 45 | R | C | 178.28 |
| | | | | 45 | R | CA | 56.82 |

| | | | | | | | |
|---|---|---|---|---|---|---|---|
| | | | | 45 | R | H | 8.49 |
| | | | | 45 | R | N | 116.06 |
| | | | | 46 | L | CA | 56.35 |
| | | | | 46 | L | H | 7.78 |
| | | | | 46 | L | N | 113.59 |
| | | | | 47 | F | C | 175.52 |
| | | | | 47 | F | CA | 58.42 |
| | | | | 47 | F | H | 7.57 |
| | | | | 47 | F | N | 112.02 |
| | | | | 48 | F | C | 175.47 |
| | | | | 48 | F | CA | 61.16 |
| | | | | 48 | F | H | 7.35 |
| | | | | 48 | F | N | 116.29 |
| | | | | 49 | K | C | 175.39 |
| | | | | 49 | K | CA | 56.98 |
| | | | | 49 | K | H | 10.00 |
| | | | | 49 | K | N | 120.63 |
| | | | | 50 | S | C | 174.77 |
| | | | | 50 | S | CA | 60.47 |
| | | | | 50 | S | H | 9.11 |
| | | | | 50 | S | N | 111.02 |
| | | | | 51 | I | CA | 65.96 |
| | | | | 51 | I | H | 9.97 |
| | | | | 51 | I | N | 126.54 |
| | | | | 52 | Y | C | 178.83 |
| | | | | 53 | R | CA | 58.93 |
| | | | | 53 | R | H | 8.56 |
| | | | | 53 | R | N | 122.28 |
| | | | | 54 | F | CA | 60.83 |
| | | | | 54 | F | H | 7.35 |
| | | | | 54 | F | N | 120.66 |

**Table S3:** Assigned chemical shifts for pore bound Rmt to $M2_{18-60}$ using 40 mM Rmt

| Bound | | | | | |
|---|---|---|---|---|---|
| residue number | Amino acid | Atom | Chain A | | Chain B |
| 26 | L | N | 118.778 | N | 118.361 |
| 26 | L | HN | 8.707 | NH | 8.455 |
| 26 | L | CA | 57.822 | CA | 58.258 |
| 26 | L | CB | 41.542 | CB | 41.45 |
| 26 | L | C | 178.685 | C | 177.989 |
| 27 | V | N | 120.4 | N | 120.608 |
| 27 | V | HN | 7.397 | NH | 7.171 |
| 27 | V | CA | 65.162 | CA | 65.397 |
| 27 | V | HA | 3.802 | HA | 3.694 |
| 27 | V | C | 180.138 | C | 180.422 |
| 28 | V | N | 121.321 | N | 123.692 |
| 28 | V | HN | 7.928 | NH | 8.222 |
| 28 | V | CA | 67.109 | CA | 67.434 |
| 28 | V | HA | 3.386 | HA | 3.316 |
| 28 | V | CB | 30.835 | CB | 31.586 |
| 28 | V | C | 178.991 | C | 178.259 |
| 29 | A | N | 123.4 | N | 120.8 |
| 29 | A | HN | 8.819 | NH | 8.598 |
| 29 | A | CA | 56.029 | CA | 56.004 |
| 29 | A | HA | 3.898 | HA | 3.772 |
| 29 | A | CB | 17.751 | CB | 17.524 |
| 29 | A | C | 179.906 | C | 179.238 |
| 30 | A | N | 119.05 | N | 118.042 |
| 30 | A | HN | 8.871 | NH | 8.787 |
| 30 | A | CA | 56.05 | CA | 55.31 |
| 30 | A | CB | 18.42 | CB | 18.546 |
| 30 | A | C | 179.896 | C | 180.086 |
| 31 | S | N | 120.186 | N | 119.462 |
| 31 | S | HN | 8.035 | NH | 8.34 |
| 31 | S | CA | 63.447 | CA | 63.67 |
| 31 | S | HA | 3.971 | HA | |
| 31 | S | CB | 61.711 | CB | 62.716 |
| 31 | S | C | 175.033 | C | 175.011 |
| 32 | I | N | 121.99 | N | 121.744 |
| 32 | I | HN | 7.603 | NH | 7.806 |
| 32 | I | CA | 66.113 | CA | 66.203 |
| 32 | I | CB | 38.16 | CB | 38.393 |
| 32 | I | C | 177.839 | C | 177.525 |
| 33 | I | N | 118.701 | N | 117.674 |
| 33 | I | HN | 8.393 | NH | 8.375 |
| 33 | I | CA | 65.689 | CA | 63.511 |

| | | | | | |
|---|---|---|---|---|---|
| 33 | I | HA | | HA | |
| 33 | I | CB | 37.377 | CB | |
| 33 | I | C | 176.646 | C | 176.84 |
| 34 | G | N | 109.328 | N | 109.451 |
| 34 | G | HN | 8.664 | NH | 8.555 |
| 34 | G | CA | 46.871 | CA | 46.855 |
| 34 | G | C | 177.438 | C | 177.284 |
| 35 | I | N | 124.219 | N | 123.39 |
| 35 | I | HN | 7.731 | NH | 7.757 |
| 35 | I | CA | 66.137 | CA | 66.143 |
| 35 | I | HA | | HA | 3.57 |
| 35 | I | CB | 37.503 | CB | 37.672 |
| 35 | I | C | 177.098 | C | 176.86 |
| 36 | L | N | 118.616 | N | 118.76 |
| 36 | L | HN | 8.053 | NH | 8.107 |
| 36 | L | CA | 57.954 | CA | 57.951 |
| 36 | L | HA | | HA | |
| 36 | L | CB | 41.583 | CB | 41.583 |
| 36 | L | C | 177.136 | C | 177.151 |
| 37 | H | N | | N | 117.772 |
| 37 | H | HN | | NH | 8.896 |
| 37 | H | CA | | CA | 57.971 |
| 37 | H | HA | | HA | |
| 37 | H | CB | | CB | 32.712 |
| 38 | L | N | | N | 116.523 |
| 38 | L | HN | | NH | 6.913 |
| 38 | L | CA | | CA | 58.133 |
| 38 | L | HA | | HA | |
| 38 | L | CB | | CB | 39.237 |
| 38 | L | C | | C | 177.637 |
| 39 | I | N | | N | 116.41 |
| 39 | I | HN | | NH | 7.496 |
| 39 | I | CA | | CA | 65.513 |
| 39 | I | C | | C | 176.633 |
| 40 | L | N | | N | 115.184 |
| 40 | L | HN | | NH | 8.584 |
| 40 | L | CA | | CA | 57.84 |
| 40 | L | HA | | HA | 3.869 |
| 40 | L | CB | | CB | 43.206 |
| 40 | L | C | | C | 177.751 |
| 41 | W | N | | N | 119.928 |
| 41 | W | HN | | NH | 8.253 |
| 41 | W | CA | | CA | 62.267 |
| 41 | W | HA | | HA | 4.196 |

| | | | | | |
|---|---|---|---|---|---|
| 41 | W | CB | | CB | 27.281 |
| 41 | W | C | | C | 177.216 |
| 42 | I | N | | N | 118.532 |
| 42 | I | HN | | NH | 8.304 |

Table S4: Normalized intensities extracted from (H)NH spectra at 25 degrees. Numbers in italic were used for extracting the slope at ~50%. The indicated error is calculated based on the spectral noise.

| KELVIN | 298.15 | | | | | | | | |
|---|---|---|---|---|---|---|---|---|---|
| TIMES (SEC) | HISTA | HISTB | GLY | ERROR HISTA | ERROR HISTB | ERROR GLY | KINETICS 50% | | |
| 0 | 1 | 1 | 1 | 0.102 | 0.171 | 0.097 | HISTA | HISTB | GLY |
| 61200 | *0.8* | *0.793* | 0.952 | 0.067 | 0.112 | 0.064 | -2.64E-06 | -2.15E-06 | -1.91E-06 |
| 126900 | 0.587 | 0.565 | *0.824* | 0.066 | 0.111 | 0.063 | ERROR MAX | | |
| 137700 | 0.547 | 0.531 | 0.771 | 0.072 | 0.121 | 0.069 | -3.66E-06 | -3.87E-06 | -2.86E-06 |
| 188100 | *0.465* | *0.52* | 0.684 | 0.063 | 0.106 | 0.06 | ERROR MIN | | |
| 251100 | 0.351 | 0.37 | *0.587* | 0.058 | 0.098 | 0.056 | -1.61E-06 | -4.31E-07 | -9.51E-07 |
| 423900 | 0.196 | 0.174 | 0.106 | 0.069 | 0.115 | 0.066 | ERROR AVERAGE | | |
| 549900 | 0.11 | 0.077 | 0.248 | 0.085 | 0.142 | 0.081 | -2.64E-06 | -2.15E-06 | -1.91E-06 |
| 629100 | 0.142 | 0.103 | 0.228 | 0.112 | 0.188 | 0.107 | | | |

Table S5: Normalized intensities extracted from (H)NH spectra at 40 degrees. Numbers in italic were used for extracting the slope at ~50%. The indicated error is calculated based on the spectral noise.

| KELVIN | 313.15 | | | | | | | | |
|---|---|---|---|---|---|---|---|---|---|
| TIMES (SEC) | HISTA | HISTB | GLY | ERROR HISTA | ERROR HISTB | ERROR GLY | KINETICS 50% | | |
| 0 | 1 | 1.00 | 1 | 0.05 | 0.06 | 0.05 | HISTA | HISTB | GLY |
| 900 | 0.959 | 0.87 | 0.969 | 0.051 | 0.06 | 0.051 | -5.34E-06 | -7.18E-06 | -5.05E-06 |
| 4500 | 1.046 | 0.99 | 0.957 | 0.047 | 0.06 | 0.047 | ERROR MAX | | |
| 11700 | 1.032 | *0.97* | *0.984* | 0.049 | 0.06 | 0.049 | -6.51E-06 | -9.97E-06 | -6.04E-06 |

| | | | | | | | | | |
|---|---|---|---|---|---|---|---|---|---|
| 26100 | *0.956* | 0.86 | 0.914 | 0.05 | 0.06 | 0.05 | ERROR MIN | | |
| 54900 | 0.683 | *0.66* | 0.804 | 0.049 | 0.06 | 0.049 | -4.17E-06 | -4.39E-06 | -4.06E-06 |
| 112500 | *0.495* | 0.40 | *0.475* | 0.051 | 0.06 | 0.051 | ERROR AVERAGE | | |
| 227700 | -0.006 | -0.04 | 0.041 | 0.053 | 0.07 | 0.053 | -5.34E-06 | -7.18E-06 | -5.05E-06 |

Table S6: Normalized intensities extracted from (H)NH spectra at 55 degrees. Numbers in italic were used for extracting the slope at ~50. The indicated error is calculated based on the spectral noise.

| KELVIN | 328.15 | | | | | | | | |
|---|---|---|---|---|---|---|---|---|---|
| TIMES (SEC) | HISTA | HISTB | GLY | ERROR HISTA | ERROR HISTB | ERROR GLY | KINETICS 50% | | |
| 0 | *1* | *1* | *1* | 0.038 | 0.06 | 0.035 | HISTA | HISTB | GLY |
| 5400 | *0.066* | *0.065* | *0.124* | 0.027 | 0.042 | 0.024 | -1.73E-04 | -1.73E-04 | -1.62E-04 |
| 9000 | 0.038 | 0.006 | 0.091 | 0.019 | 0.029 | 0.017 | ERROR MAX | | |
| | | | | | | | -1.85E-04 | -1.92E-04 | -1.73E-04 |
| | | | | | | | ERROR MIN | | |
| | | | | | | | -1.61E-04 | -1.54E-04 | -1.51E-04 |
| | | | | | | | ERROR AVERAGE | | |
| | | | | | | | -1.73E-04 | -1.73E-04 | -1.62E-04 |

Table S7: Values obtained from the kinetics that were used for the Arrhenius analysis. The average was calculated including the error at each point and extracting maximal and minimal slope. The error is calculated using the standard deviation of the three values obtained for each temperature (average slope, Maximal slope and minimal slope).

| 10E03/T | HistA | HistB | Gly | Average |
|---|---|---|---|---|
| 3.354 | -2.64E-06 | -2.15E-06 | -1.91E-06 | -2.23E-06 |
| 3.193 | -5.34E-06 | -7.18E-06 | -5.05E-06 | -5.86E-06 |
| 3.047 | -1.73E-04 | -1.73E-04 | -1.62E-04 | -1.69E-04 |

| | | | | |
|---|---|---|---|---|
| **3.354** | -12.845 | -13.050 | -13.171 | -13.022 |
| **3.193** | -12.140 | -11.844 | -12.196 | -12.060 |
| **3.047** | -8.662 | -8.662 | -8.728 | -8.684 |
| **Slope** | -13.642 | -14.308 | -14.488 | -14.146 |
| **Ea** | -113.416 | -118.960 | -120.457 | -117.611 |
| **Average** | -114.850 | -128.079 | -123.025 | -121.985 |
| **Error** | 9.240 | 26.915 | 13.114 | 16.423 |

Table S8. Acquisition parameters for (H)NH spectra recorded for tracking of kinetics. Spectra were recorded at 18.8 Tesla in a triple channel 1.3 mm probe with the parameters indicated.

| | $^{1}$H | | $^{15}$N | |
|---|---|---|---|---|
| | **Time (ms)** | **Number of points** | **Time (ms)** | **Number of points** |
| DPhPC | 21 | 1024 | 8 | 156 |
| DPhPC + Chol | 21 | 1024 | 8 | 156 |
| VM | 21 | 1024 | 8 | 156 |
